# Secret life of prophages: template-directed synthesis of DNA superstructures via prophage activation and rolling circle replication in bacterial biofilms

**DOI:** 10.64898/2025.12.02.691978

**Authors:** Gabriel Antonio S. Minero, Line Mørkholt Lund, Lasse Hyldgaard Klausen, Obinna Markraphael Ajunwa, Mingdong Dong, Rikke Louise Meyer, Victoria Birkedal, Kai Thormann

## Abstract

Extracellular DNA (eDNA) plays crucial roles in biofilm formation and function, yet the role of bacteriophages (phages) in controlling eDNA synthesis, structure and activity remains obscure. Here, we demonstrate that phages harbored by environmental bacteria can be exploited for programmable synthesis of functional eDNA superstructures. We designed a 112-nucleotide circular template (T1) and used it to direct rolling circle replication (RCR) of G-quadruplex (GQ) motifs in *Shewanella oneidensis* and *Bacillus subtilis*. Under nutrient-limiting conditions, prophage activation triggered cell lysis and subsequent extracellular DNA synthesis, producing multimeric GQ concatemers that self-assembled into distinct morphologies: spherical structures (≤10 μm) in *S. oneidensis* and wire-like structures (>50 μm) in *B. subtilis*. Real-time monitoring using fluorescent reporter strains revealed that DNA synthesis occurred predominantly after bacterial lysis, coinciding with prophage replication. The resulting DNA superstructures exhibited peroxidase activity through GQ-hemin DNAzyme formation and enhanced the electrochemical properties of *S. oneidensis* biofilms, showing a 3-fold increase in current density. This work unveils a previously unknown mechanism by which prophages contribute to biofilm architecture and establishes a biotechnological platform for engineering functional DNA materials in living bacterial communities, with potential applications in biotechnology and synthetic biology.

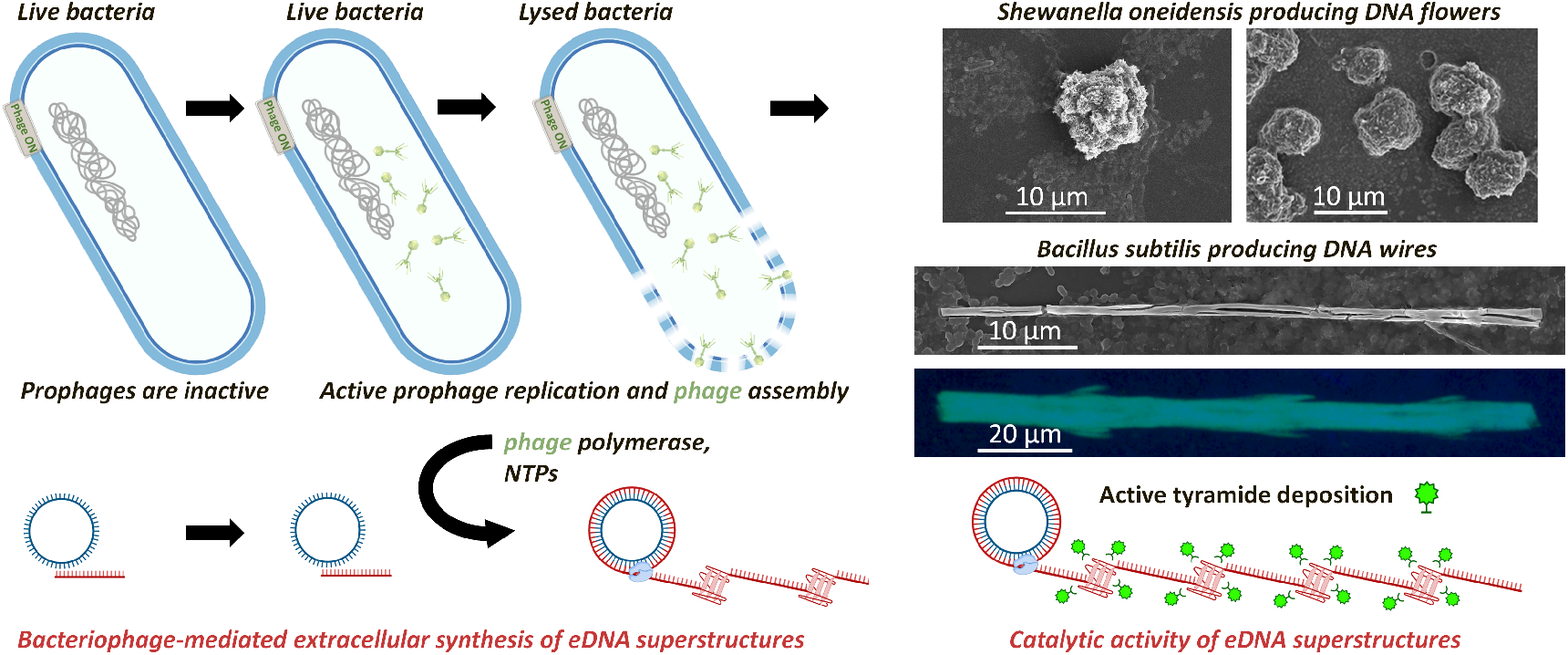

## INTRODUCTION

Bacteria release and harness nucleic acids as structural materials in biofilms - microbial communities encased in self-produced polymeric matrices. While extracellular DNA (eDNA) is known to provide structural support for the biofilm’s extracellular matrix, recent research also points to eDNA as a catalyst and a conduit for extracellular electron transport (EET). The electron transfer is facilitated by eDNA’s ability to bind redox-active molecules like phenazines (1) or hemin (2), allowing bacteria to respire in the absence of electron acceptors by ferrying electrons to an electrode or to oxygen located at some distance from the cell. The unique structural properties of non-canonical DNA forms play a key role in eDNA-mediated EET (2). G-quadruplexes (GQ) were shown to scaffold bacterial redox-active molecules such as cytochromes, maintaining them in spatial arrangement for a rapid electron transfer (3). GQs facilitate π-π interactions and bind hemin with high specificity, creating a GQ/hemin DNAzyme with peroxidase-like activity (4). The DNAzyme activity provides an additional mode for electron transfer from bacterial metabolic processes to hydrogen peroxide and potentially other electron acceptors, which could catalyze CO_2_ reduction (5,6) or degrade micropollutants (7). EET mediated by GQ/hemin may therefore augment microbial processes in biofilms used in e.g. *Shewanella oneidensis* (*S. oneidensis*) microbial fuel cells (8) or activated sludge (9) to tackle environmental challenges. Inducing catalytic DNA production to augment these processes require knowledge about how they form, and their formation can be stimulated in the complex microbial communities that carry out these processes.

In addition to bacteria, biofilms also contain bacterial viruses (bacteriophages). These phages can facilitate eDNA release from cells into the biofilm’s extracellular matrix (10-12). Globally, phages outnumber bacteria tenfold (13,14). Virulent phages infect and replicate inside bacterial cells, causing cell lysis and the extracellular release of hundreds of new phage virions (15). In contrast, temperate phages can lie dormant in bacterial genomes as prophages, which are activated by certain conditions that trigger the SOS response to oxidative stress and DNA damage, e.g. in the presence of mitomycin C (15,16) or iron (10). The SOS response can also be induced in cells post starvation, i.e. upon reintroduction of carbon (17). Prophages are present across a variety of bacterial species (14,18) and the mechanisms for prophage activation vary (19). Some prophages, e.g. Mu, activate replication in response to starvation (20) whereas prophage λ remains in its lysogenic state under cell starvation (21,22). In *Bacillus subtilis* (*B. subtilis*), most predicted prophages activate spontaneously during starvation-induced sporulation (23). Therefore, starvation-induced SOS response can be utilized as a generic approach to activate some prophages in environmental species and to stimulate eDNA release along with progeny phages.

Some phages, including prophages λSo/ MuSo in *S. oneidensis* and SPP/ SPβ in *B. subtilis*, contain unique DNA polymerases that can produce ssDNA from circular ssDNA templates by rolling circle replication (RCR) (24-26). Phage polymerases can thus be used to synthesize large quantities of multimeric DNA cocoons *in vitro* from circular templates (27). If unlimited by nucleotide triphosphates (NTPs), circular DNA replication can produce linear concatemer products of thousands of “stitched” complementary copies of the circular DNA template. These products can even be programmed to include multimeric GQ-DNA (28). Furthermore, the incorporation of NTPs into growing DNA chains releases pyrophosphate (PP) anions as PP precipitates. These insoluble PPs can then nucleate the formation and phase separation of micrometer-sized flower-like DNA superstructures in cell-free systems (29,30). The size and morphology of these “DNA flowers” is unique and depends on the sequence and concentration of the DNA template, the type of divalent cation and even the type of the polymerase (31). DNA sequence, structure, and supramolecular organization are therefore all intimately connected. While this phenomenon has been demonstrated *in vitro*, it has never been reproduced in bacteria or their biofilms.

We hypothesized that phage polymerases can continue to synthesize DNA in the biofilm matrix after they are released by cell lysis, where the crowded microenvironment around lysed bacteria in biofilms still contain polymerases and NTPs. Subsequently, sequence-specific DNA structures and topologies can be synthesized and consequently integrated into bacterial eDNA matrices, by promoting extracellular DNA replication using the right exogeneous template. Considering the structural importance of eDNA and the other key eDNA functions e.g. eDNA mediated EET (1,2), incorporating specific DNA motifs into the biofilm matrix can enhance biofilm’s form and function. The aim of this study is therefore to explore synthesis, shapes and functions of DNA superstructures by template-directed synthesis of functional DNA structures in the extracellular matrix of biofilms from the environmental microorganisms *S. oneidensis* and *B. subtilis*. The outcomes of this study will enable enhanced performance of key environmental bioprocesses using, e.g., electroactive *S. oneidensis* microbial fuel cells (8) and methanogenic archaeon (6).

## RESULTS

### Extracellular DNA synthesis is possible after bacterial lysis by phage activation

We sought to establish if extracellular DNA replication can occur in planktonic cultures where NTPs, phage polymerase and other cellular materials are released by phage-induced cell lysis. We used a circular ssDNA template T1 (Table 1) generating a GQ concatemer comprised of several hundred stacked G-tetrads stabilized by Hoogsteen hydrogen bonds between the guanines holding G-tetrads in a square planar configuration (32). To detect RCR, we used GQ-specific N-Methyl Protoporphyrin (NMP) dye (33). The RCR kinetic plots were obtained by subtracting NMP fluorescence in the non-templated cultures from the NMP fluorescence in the templated cultures resulting in the “NMP difference” value at every time point (Figures 1+S1). Extracellular DNA synthesis by RCR was indicated by an increase in the NMP difference, describing the difference in NMP fluorescence with the non-templated culture. NMP binds specifically to GQ, thus GQ-DNA synthesis by *S. oneidensis* MR-1, S2391, and S5558 cultures commenced after a 300 min lag phase (Figure 1, red). At the same time, cyan fluorescent protein (CFP), representing living intact bacteria, decreased over time starting from about 200 min (Figure 1B-C, violet), indicating that bacteria were lysing due to phage activation. The Venus fluorescence (indicative of phage activation) peaked around 300 min (Figure 1B-C, green) and the signal of green fluorescent protein (GFP) fused to λSo capsid (indicative of phage assembly) peaked at 400 min (Figure 1D, yellow green). GQ-DNA therefore accumulates in response to phage activation and lysis of the *S. oneidensis* wild-type and ΔMuSoI ΔMuSoII. Phage polymerase is ultimately responsible for GQ-DNA replication (Figure 1: red curves). On the contrary, there was no increase in NMP difference in the S6610 (ΔλSo ΔMuSoI ΔMuSoII) strain lacking any phages (Figure 1B-C: blue curves). The timing of GQ-DNA synthesis coincided with activation of prophages detected in the reporter strains of *S. oneidensis* (around 300 min in the starved cultures), supporting our hypothesis that RCR from our template can occur extracellularly following phage-induced lysis of *S. oneidensis* (Figure 1B-C green and red curves indicative of phage activation and GQ synthesis, respectively; Figure 1D chartreuse curve indicative of phage λSo capsid production). The averages ± SD for the NMP difference in response to RCR facilitated by wild-type *S. oneidensis* under nutrient-limiting conditions (starvation) are shown in Figure 2A.

**Table 1.**
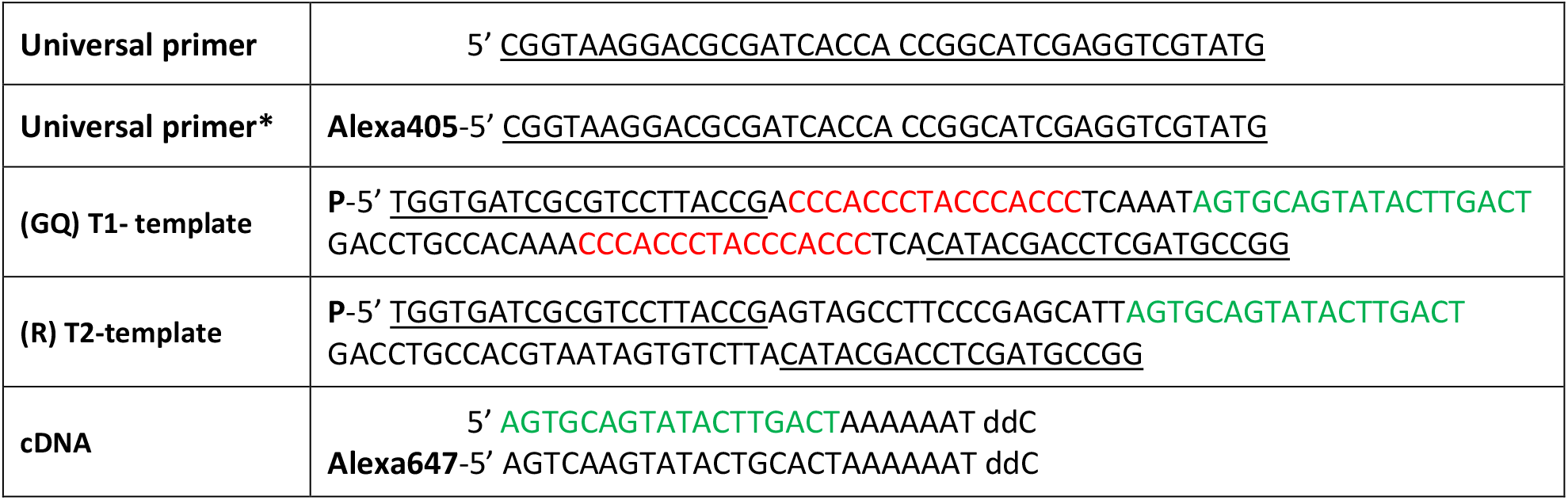
The oligonucleotides purchased from IDT and used for the synthetic ligation of circular DNA. Underlined are complementary regions between DNA “Universal primer” and “Templates”. In red are the motifs encoding for GQ-DNA synthesis. The circularizing oligonucleotides were phosphorylated at the 5’-end. A complementary DNA oligonucleotide (cDNA) contained 3’-dideoxycytosine and 5’-Alexa647. The sequence of cDNA is highlighted in green.

**Figure 1.**
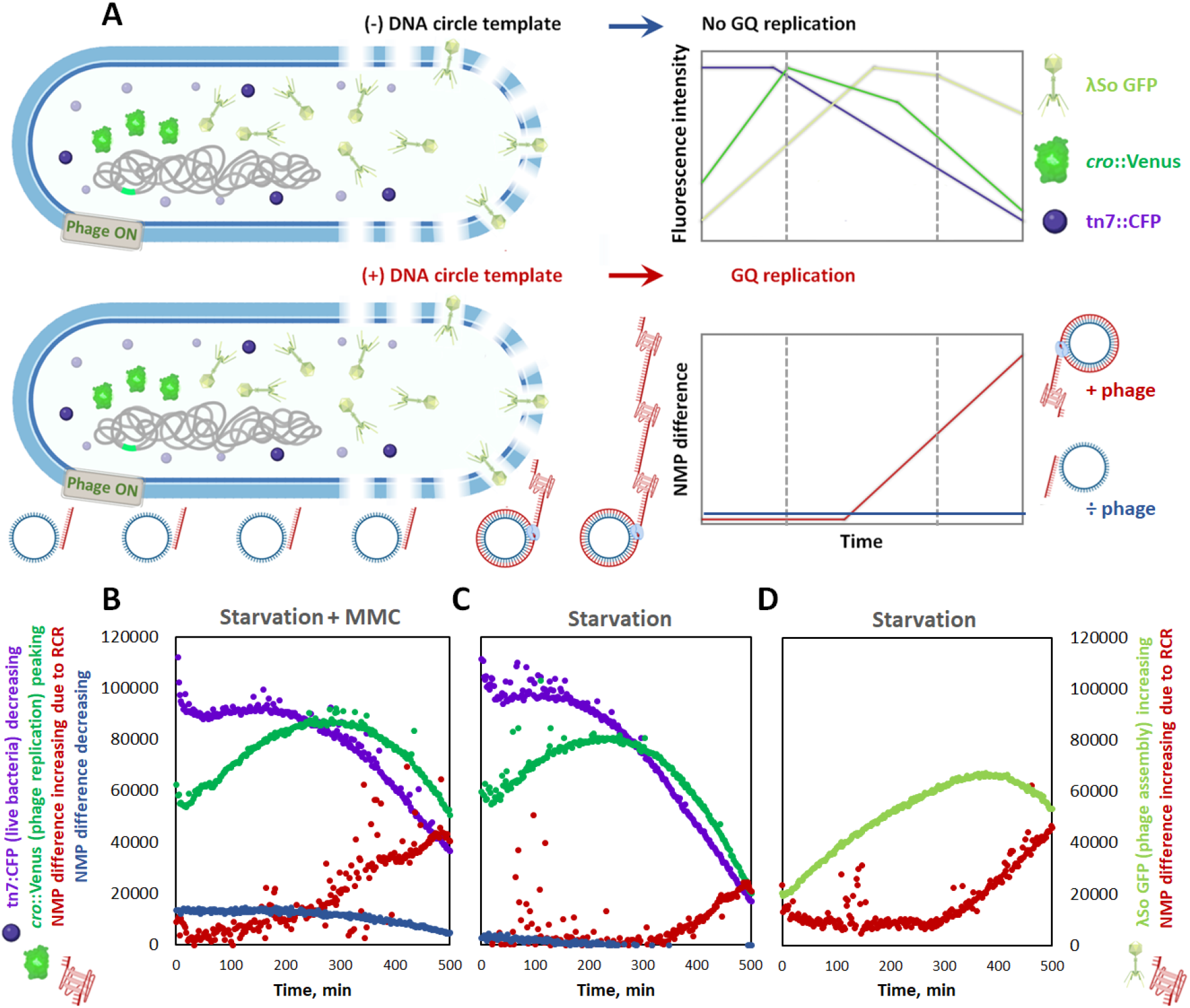
Time-resolved phage proliferation and bacterial lysis lead to RCR and synthesis of multimeric G-quadruplex DNA from an exogenous template. (A) Conceptual presentation of a *S. oneidensis* cell in time. *S. oneidensis* MR-1 cells were harvested from overnight cultures in LB media. In the beginning, bacteria are intact. Upon phage proliferation and assembly, bacteria lyse and lead to extracellular replication of GQ-DNA in the vicinity of the lysed cell. Schematic representation of phage replication detected by chromosomal *cro*::Venus fusion (the protein as well as its promoter, are shown in green) in the wild-type S2391 (live bacteria expressing *tn7*::eCFP shown in violet) and phage assembly (yellow green) detected by GFP fused to λ-capsid in the S5558 (ΔMuSoI ΔMuSoII). Replication of the GQ-DNA in the presence of the circular DNA T1 is detected by NMP. “NMP difference” (shown in red) was obtained after deducting the NMP signal of the control (i.e., non-templated) cultures. The NMP difference obtained in the wild-type (+ phage) and mutant (ΔλSo ΔMuSoI ΔMuSoII) cultures is shown in red and blue, respectively. (B-D) Real-time monitoring GQ-DNA via NMP difference and phage proliferation via the fluorescent proteins expressed by the reporter strains. To activate phage replication in the wild-type S2391 (*cro*::Venus fusion) reporter strain, we used mitomycin C in (B) and starvation in (C). To activate replication of phage λSo and assembly of λSo-GFP capsids in the S5558 reporter strain, we used starvation in (D).

**Figure 2.**
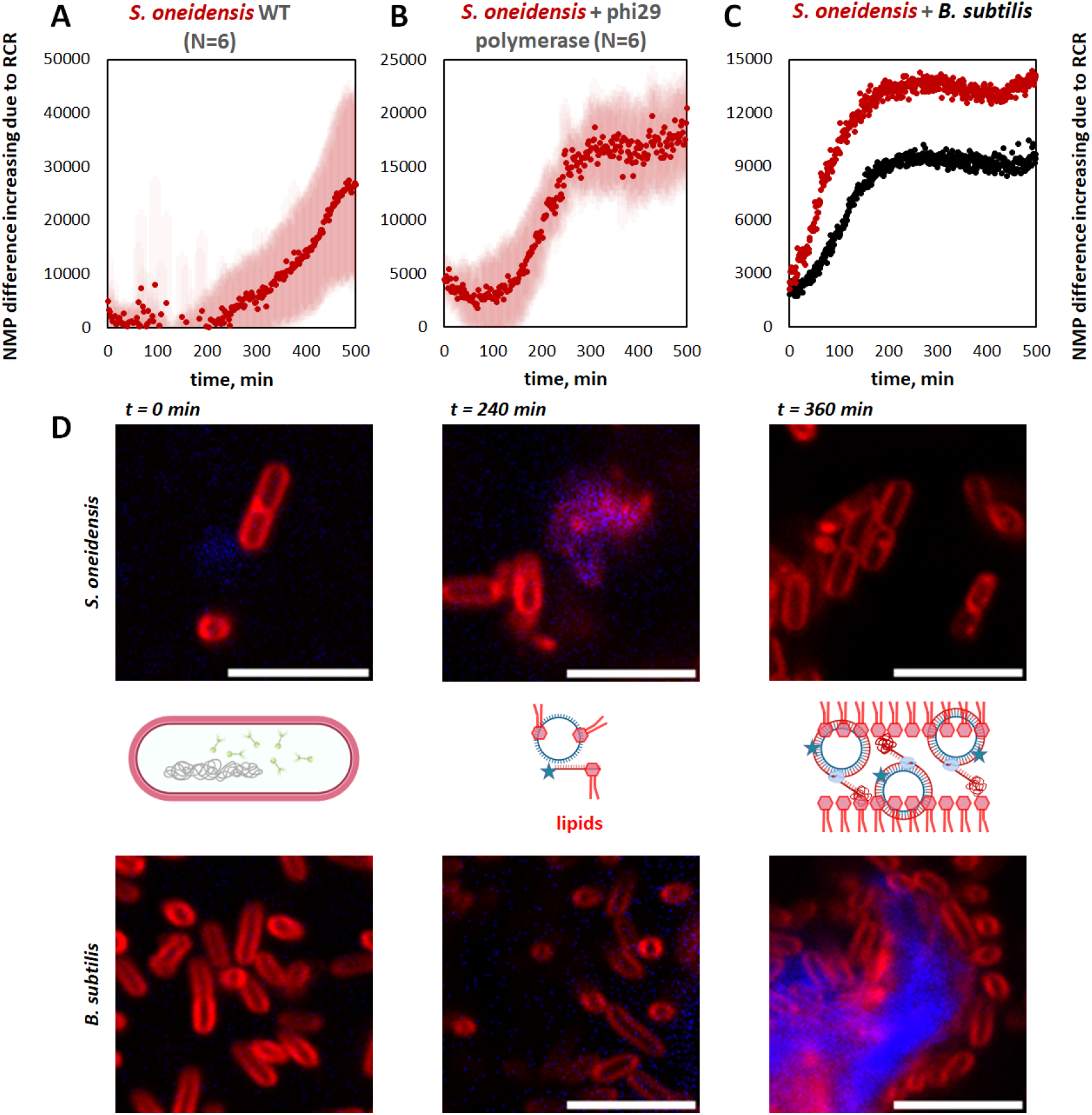
DNA replication and assembly of superstructures is detected as “NMP difference” in planktonic cultures of *S. oneidensis* and *B. subtilis*. (A) *S. oneidensis* wild-type strains. (B) *S. oneidensis* wild-type and ΔMuSoI ΔMuSoII strains supplemented with phi29 polymerase (20 u). The NMP difference is shown after deducting controls without polymerase. (C) *B. subtilis* monoculture (black) and *B. subtilis* co-cultured with *S. oneidensis* (red), shown as averages of two biological replicates. (D) Fluorescence imaging with single photon counting confocal microscopy of FM 4-64 lipophilic membrane stain (in red) and Alexa405-DNA circle (in blue) detected at time 0 and after 240 and 360 min of continuous incubation of bacterial cells with the DNA. Scale bar 5 μm. In the middle is the proposed scheme of DNA templates being trapped in the phospholipids and replicated in the extracellular space. See Figure 1 for the proposed mechanism of RCR.

To determine whether DNA replication could have occurred intracellularly following uptake of the DNA template, we added exogenous phi29 polymerase (Thermofisher scientific), which cannot be taken up by the bacteria, to the *S. oneidensis* cultures in the buffer containing the circular DNA template. Addition of phi29 resulted in an earlier increase in NMP fluorescence (at *t* > 100 min, Figure 2B).

To corroborate our results, we performed a similar experiment in *Bacillus subtilis*, another environmental species that harbors several phages that use RCR: SPP1 (34), SPβ (35), phi3T (23), phi105 (36) and phi29 (37). The latter encodes phi29 polymerase, which is very robust in processivity (30) and which we used as a booster of extracellular RCR (Figure 2B). Figure 2C shows an immediate increase in GQ-DNA synthesis (quantified by the “NMP difference”) in *B. subtilis* monoculture and in *B. subtilis + S. oneidensis* co-culture. This suggests that *B. subtilis* phage replication machinery can also facilitate extracellular GQ-DNA synthesis from the circular template amended to the culture, and together these results support our hypothesis that RCR can occur in the extracellular space following the onset of starvation.

Finally, to test a possible uptake of the circular DNA by bacterial cells, we employed a fluorescently labelled universal primer (Table 1) and the T2 template, resulting in the synthesis of R-DNA. Imaging of the uptake of the DNA circle by *S. oneidensis* and *B. subtilis* detected the fluorescent DNA (blue) associated with clumped deformed membranes of the bacteria at *t* > 240 min, suggesting interaction of lysed bacterial membranes with the DNA circle, creating a crowded environment for subsequent extracellular DNA replication via RCR. Finally, at *t* > 360 min, we detected larger superstructures (only in *B. subtilis*) resembling “wires”. We could not detect an uptake of the fluorescent DNA by either *S. oneidensis* or *B. subtilis* at *t* < 360 min (Figure 2D). Collectively, our results using planktonic cultures and NMP fluorescence to detect GQ-DNA synthesis in real-time suggest that both *S. oneidensis* and *B. subtilis* can replicate an exogenous circular DNA template after prophage activation under starvation, and this replication may take place in the extracellular environment.

### Phage-mediated DNA synthesis in *S. oneidensis* biofilm results in spherical DNA superstructures

RCR is known to produce a DNA concatemer assembled into a superstructure of DNA and various insoluble pyrophosphates (29), which are often described as DNA flowers and can reach several microns in size. Our results indicate production of GQ-DNA in planktonic *S. oneidensis* cultures, and we hypothesized that concatemers of GQ-DNA could form superstructures with other biopolymers in *S. oneidensis* biofilms that experience a higher degree of molecular crowding than planktonic cultures.

To test this hypothesis, we inoculated *S. oneidensis* biofilms from the planktonic cultures starved in the buffer and transferred it to TSB media, where the surviving bacteria were cultivated for additional 2 days to form a biofilm with GQ-DNA integrated into the extracellular matrix. eDNA was visualized by confocal laser scanning microscopy (CLSM) using fluorescence immunolabelling with the GQ-specific antibody BG4 (4), the live/dead staining by DNA-binding dyes SYTO™60/TOTO™-1 and the gold standard GQ-staining by NMP (33). Signals from BG4, TOTO™-1 and NMP staining revealed that GQ-DNA was located inside extracellular spherical superstructures (Figure 3). Intriguingly, the DNA-binding dyes SYTO™60 and TOTO™-1 promoted Förster Resonance Energy Transfer (FRET) within these spherical superstructures (yet not inside either live or dead cells that were stained by SYTO™60 and TOTO™-1, respectively, Figure 3B). This phenomenon is linked to simultaneous binding of both dyes and excitation of TOTO ™-1 (donor), which energy is then transferred to SYTO™60 (acceptor) leading to detection of SYTO™60 emission. The acceptor emission, i.e. FRET, is likely to be GQ-specific (38) due to the compact “barrel” shape of GQ, while donor, i.e. TOTO™-1, emission is specific for B-DNA (ESI, Figure S2). Moreover, TOTO™-1 is cell impermeable and, therefore, FRET signal is specific for eDNA whereas NMP is cell-permeable (Figure 3C). Ultimately, we claim detection of GQ motifs in the spherical structures of up to 10 μm in size and what appeared as extracellular aggregates of these spheres using three independent GQ-specific staining assays (Figure 3A-D). Neither of the non-templated *S. oneidensis* WT biofilms (Figures S3) or templated *S. oneidensis* MUT (ΔλSo ΔMuSoI ΔMuSoII) biofilms lacking prophages (Figure S4) produced the spheres, confirming that phage replication and the externally added ssDNA circular template are responsible for the synthesis of GQ-rich DNA and the formation of the superstructures. The FRET method and additional probes confirmed the extracellular nature of the synthesized DNA (Figure S2). The structures were identified to be organic, also containing sodium, potassium and phosphorous (Figure 3F).

**Figure 3.**
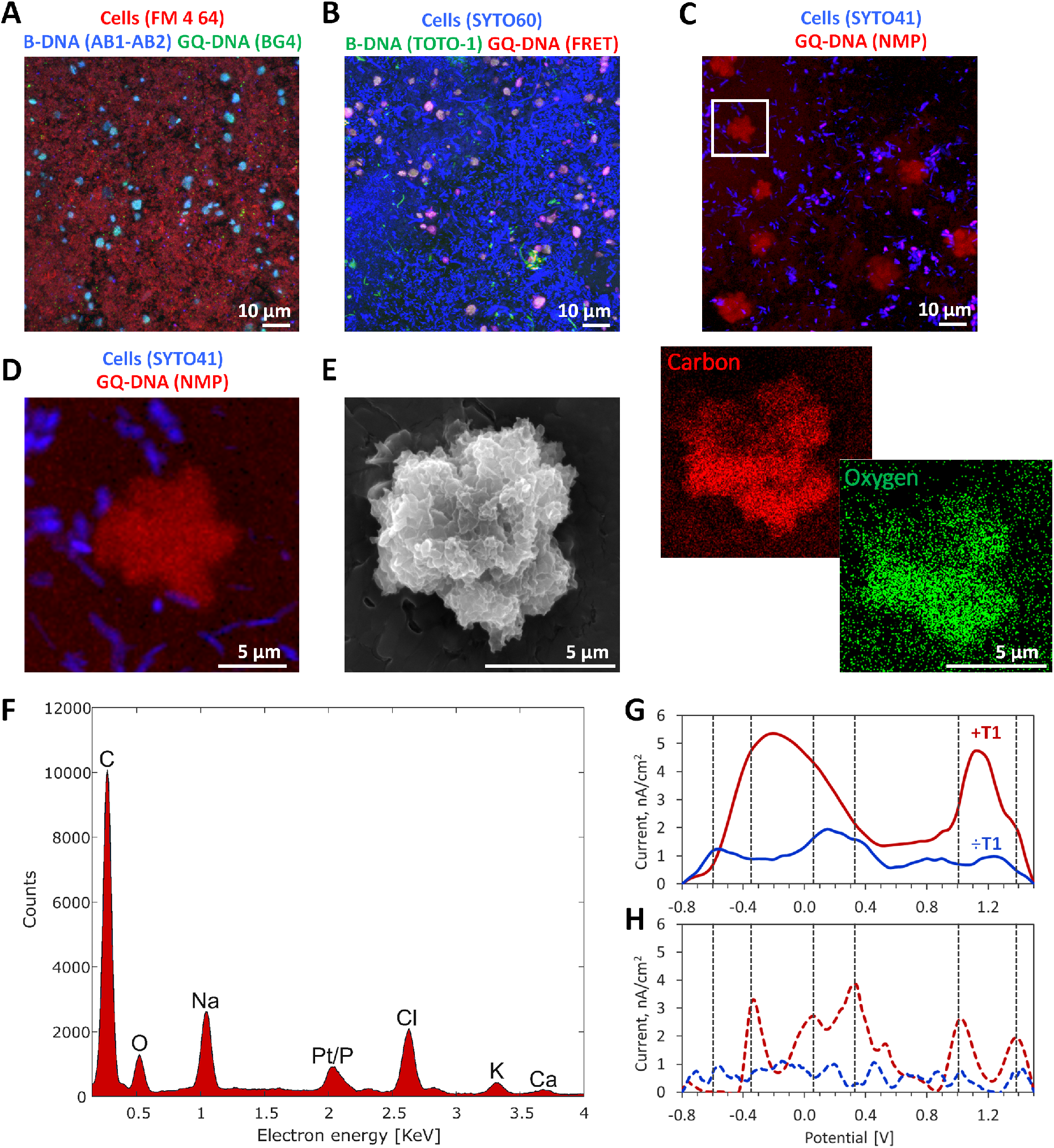
Replication of the circular DNA T1 in *S. oneidensis* biofilms leads to formation of GQ-rich crystal-like spherical superstructures. 3D CLSM images of the biofilms stained with (A) BG4 antibody (green), (B) DNA-binding dyes TOTO™-1/ SYTO™60 resulting in FRET (red) and (C-D) GQ-specific NMP dye (red). See Figure S2 for further details of the FRET method. (E)-(F) SEM-EDX elemental analysis of the GQ-DNA crystal-like superstructure. (G-H) DPV analysis of redox activity of hemin and other redox-mediators in the wild-type *S. oneidensis* T1-templated (red curve) and non-templated (blue curve) biofilms. DPV analyses were performed on (G) 3-day biofilms under aerobic and (H) 6-day biofilms under anaerobic conditions. The templated *S. oneidensis* biofilms were doped with the DNA template T1 (150 nM) added exogenously to the TSB-NaCl media.

So far, our results show that we can prompt bacteria to produce GQ-rich DNA superstructures that are synthesized extracellularly and integrated into the extracellular matrix of biofilms. The catalytic properties of these structures could impact redox reactions and EET in the biofilm and potentially augment the activity of *S. oneidensis* biofilms for e.g. microbial fuel cells. We therefore proceeded to quantify the impact of the DNA superstructures on the redox activity of DNA templated and non-templated *S. oneidensis* biofilms grown aerobically on screen printed graphite electrodes, with the biofilms containing 0 and 150 nM T1 template.

The DPV measurements (applied in the voltage range from −0.8 to 1.5 V) detected maximum peak currents densities of approximately 1.8 nA/cm^2^ in a non-templated biofilm (Figure 3G, blue curve), and a 3-fold increase in the current density (approximately 5.3 nA/cm^2^) in the templated biofilm (Figure 3G, red curve). The influence of redox activities in the two biofilm forms was observed as multiple distinct asymmetrical peaks. In the non-templated biofilm, the peaks spanned potential from −0.6V to 1.3V. In the templated biofilms, two distinct peaks had broad baselines spanning negative to positive potentials from −0.5V to 0.4V and from 0.9V to 1.4V with midpoints of −0.3V and 1.15V, respectively. We believe that these peaks correspond to convoluted redox reactions based on their broad redox potential reaching up to 900 mV, and that the GQ-rich eDNA superstructures impacted these redox processes by scaffolding redox mediators such as hemin (2), cytochromes (3) and flavins (39).

In the aerobic form of growth, bacteria are not forced to use EET and therefore may be interpreted as not “connected” to the electrode. However, the DPV measurements suggest that the spherical superstructures are forming the connective component that can conduct electrons leading to convoluted redox processes. To omit influence of oxygen onto the redox processes, we tested these biofilms grown for additional 3-days under anaerobic conditions (Figure 3H). Again, the templated *S. oneidensis* biofilms contained several peaks in the DPV diagram which were not present in samples of non-templated *S. oneidensis* biofilms, indicating that DNA superstructures are redox-active (Figure 3H). The DPV profile was different from the initial 3-day of aerobic biofilm growth, suggesting a shift in metabolic processes over time. Although we did not investigate specific molecules eliciting the measured redox properties, we hypothesize that the overall changes in the redox activity could be due to the presence of secreted redox mediators trapped in DNA superstructures in the templated biofilms. It is important to note that the magnitude of current densities observed in this study could have been reduced by the aerobic culture method applied here to initially form these superstructures.

Our results strongly indicate that templated RCR occurred extracellularly, but we cannot exclude that some bacterial cells could take up the template and initiate synthesis of the GQ-forming concatemer inside the cell. To address if intracellular synthesis from the circular template is possible, and to visualize what it looks like, we prompted uptake of the template by adding it to the bacterial suspension and exposing the bacteria to a heat-shock treatment, which is commonly used to induce transformation in Gram negative bacteria. In *E. coli*, a rapid change of temperature from 42° to 0°C in the presence of Ca^2+^ (known as “heat-shock”) was shown to alter lipid structure leading to uptake of DNA (40). We wondered if our circular DNA can be taken up by *S. oneidensis* cells “heat-shocked” in the buffer used to induce RCR. In that case, we wondered if we could trace intracellular synthesis of GQ-DNA from DNA template T1 hijacking the phage polymerase earlier (before the lysis) and, therefore, potentially altering the phenotype of the resulting superstructures. To test this research question, we explored replication of the circular DNA in the wild-type *S. oneidensis* cells pre-treated at 42°C (2 min) and cooled at 5°C (1 h). Figure 4 shows biofilm formed by such “heat-shocked” bacterial cells stained by the intracellular SYTO™60 and extracellular TOTO™-1 dyes, resulting in the GQ-specific FRET effect (shown in red). We detected SYTO™60 and FRET signals inside bacteria (Figure 4A; Figure S2) that looked different from the spherical superstructures in Figure 3A-C as they had a lipid barrier. The FRET signal was clearly confined within bacteria, and the bacteria became enlarged deformed and permeable for TOTO™-1 dye. SEM detected the presence of crystal-like conductive biopolymers inside and outside bacterial cells (Figure 4B). The larger superstructures (resembling “X-shapes”) predominantly exhibited TOTO™-1 signal (green, Figure 4A) that is known to light up (i.e. to become detectable) when bound to the canonical B-DNA. This could be for example phage λSo double-stranded DNA. Ultimately, the weight of folded GQ-DNA may be lower within the “X-shapes” compared to the spherical superstructures. The elemental analysis of the precipitated GQ-DNA resulting from the “intracellular” replication (Figure 4C) was similar to the spherical superstructures resulting from the “extracellular” replication of GQ-DNA (Figure 3F). From these experiments, we concluded that “heat-shocked” *S. oneidensis* in the presence of the T1 template produced enlarged and deformed phenotype of cells indicating uptake of the circular template and the intracellular mode of GQ replication.

**Figure 4.**
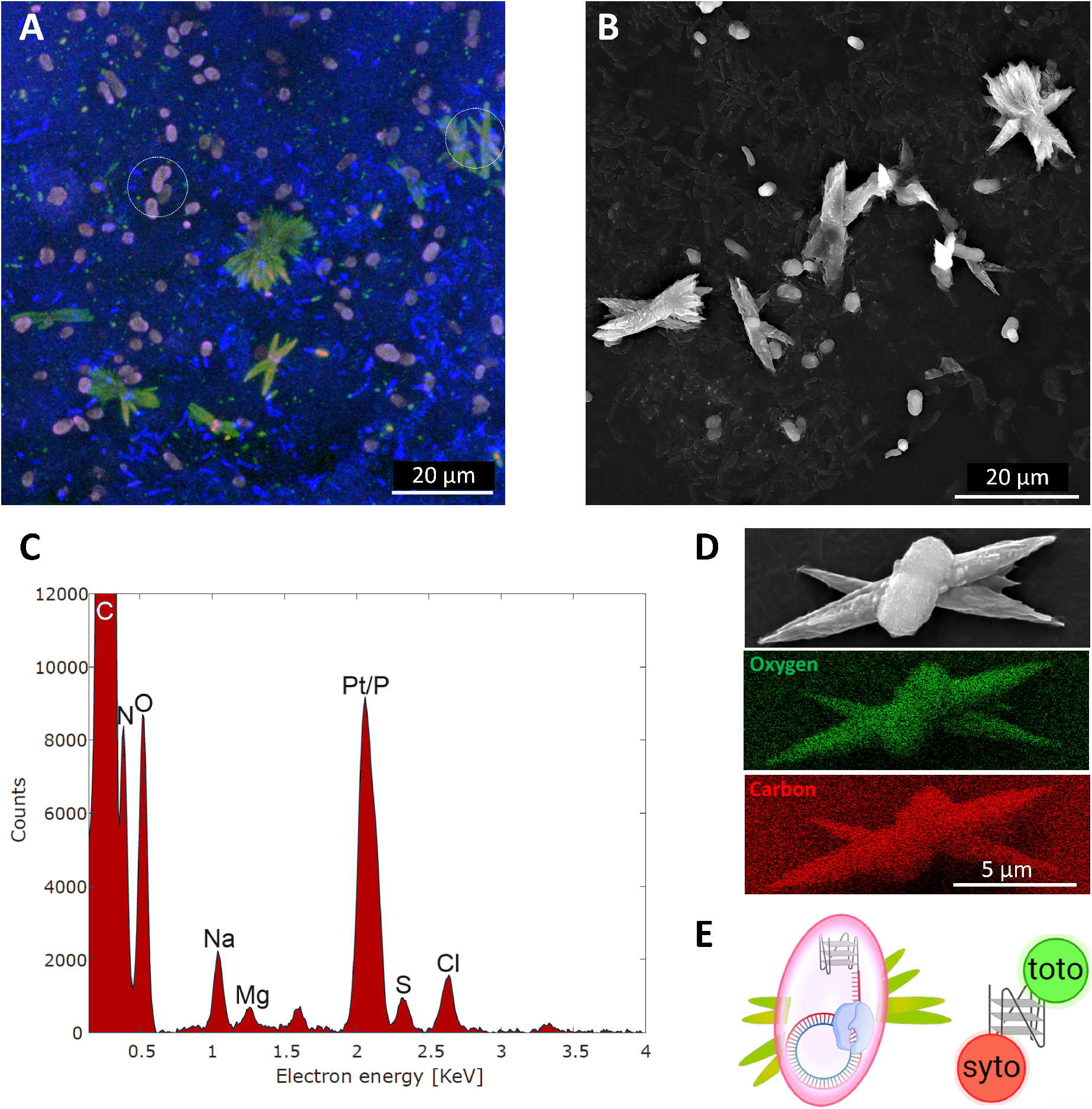
Intracellular DNA synthesis and extracellular shapes formed in DNA-templated monospecies *S. oneidensis* MR-1 in TSB-KCl, imaged by (A) CLSM, (B) SEM, and (C-D) SEM-EDX. (E) The cartoon shows bacterial cells from (A), indicated by the white circles, stained by the DNA-binding dyes SYTO™60 (acceptor excitation, acceptor emission in blue) and TOTO™-1 (donor excitation, donor emission in green) resulting in the FRET (donor excitation, acceptor emission in red). The difference from Figure 3 is the phenotype of the resulting templated biofilms, in which FRET-stained DNA products of RCR appear to be inside enlarged and deformed bacteria. See Figure S2 for further details of the FRET method.

### Phage-mediated DNA synthesis in *B. subtilis* biofilm results in wire-like superstructures

We have now provided proof-of-concept for extracellular DNA synthesis and crystallization of redox-active DNA superstructures in *S. oneidensis* biofilms. In principle, this could be a generic concept that can be applied in other microorganisms that encode prophages with polymerases capable of RCR. To demonstrate the concept in another species, we showed that *B. subtilis* encoding prophages SPP/SPβ followed a similar but faster kinetics of RCR detected by the “NMP difference” (Figure 2C), indicating synthesis of GQ-DNA. Next, we wondered if *B. subtilis* also produced GQ-DNA superstructures in biofilms. Using starved *B. subtilis* to inoculate biofilms, we observed elongated wire-like superstructures in templated biofilms. In 3-day biofilms, wires occurred in both templated and non-templated biofilms, which could be result of phage activity and replication of phage and spurious DNA. Therefore, these wire-like superstructures could contain phage or genomic DNA in addition to the programmed GQ-sequence. To ultimately differentiate the products of the templated RCR from other DNA replication, we detected peroxidase activity of the GQ-DNAzymes (41) synthesized from T1 using fluorescent tyramide signal amplification (TSA) (Figure 5A, in green). In this assay, hemin bound to GQ motifs catalyzes reduction of hydrogen peroxide and covalent addition of the tyramide residues to eDNA and other macromolecules in the vicinity of the catalyst. Furthermore, we detected binding of a fluorescently labelled oligonucleotide cDNA (Figure 5B, in blue) that is complementary to the DNA chain synthesized via RCR from either T1 or T2 template (Table 1). Figure 5 shows a DNA “wire” produced in *B. subtilis* biofilm containing both the tyramide and cDNA signal and visualized using CLSM and SEM.

**Figure 5.**
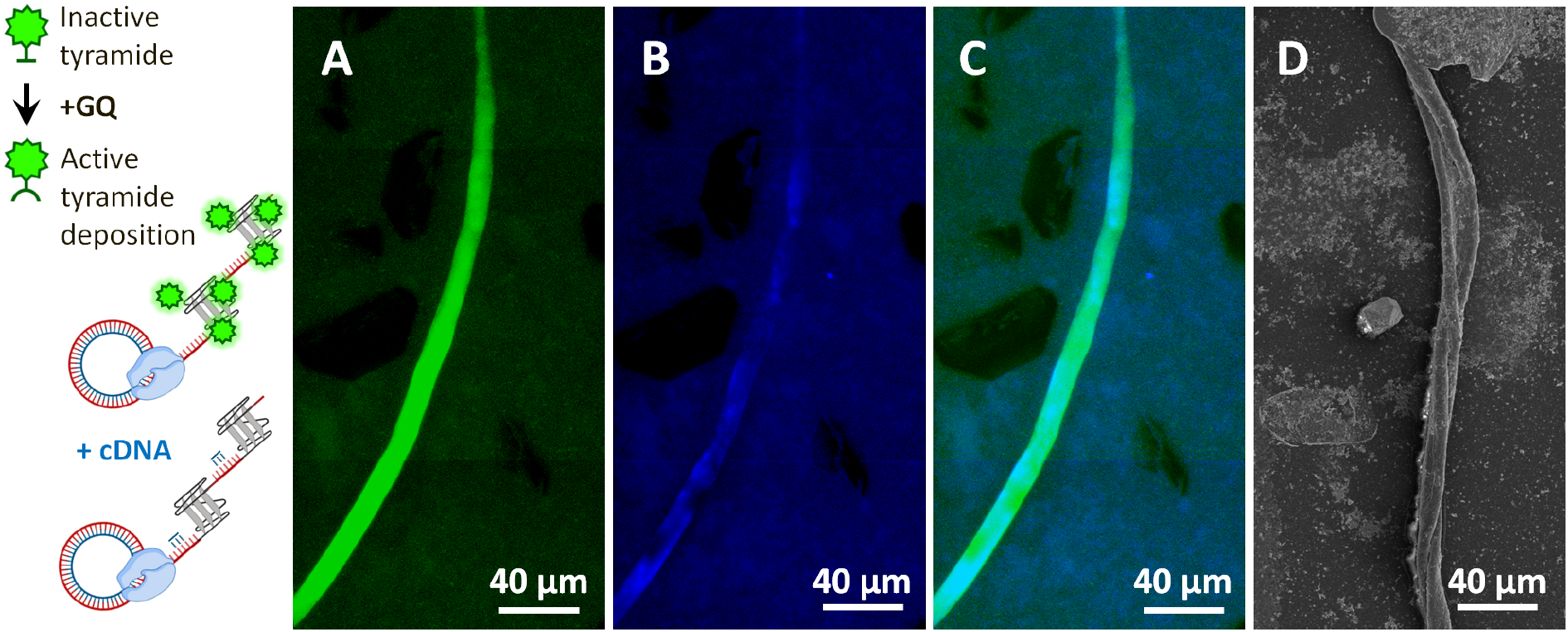
Replication of the circular DNA in *B. subtilis* biofilms leads to formation of crystal-like “wires”. Schematics show DNAzyme-mediated deposition of fluorescent tyramide (green) in the vicinity of the GQ-DNAzymes followed by hybridization and retention of fluorescent cDNA (blue). GQ-DNA superstructures were synthesized in 3-day *B. subtilis* biofilms in TSB-NaCl with exogenous T1 (150 nM), simultaneously labelled by (A) tyramide residues and (B) cDNA and visualized by CLSM. (C) Merged CLSM image and (D) SEM image of a twisting DNA wire.

### *B. subtilis* forms DNA “wire” superstructures in multispecies biofilms

*B. subtilis* as well as *S. oneidensis* are environmental strains that are being harnessed for applications such as wastewater treatment, CO_2_ capture, and bioenergy production (42,43). Both strains triggered replication of DNA from a template provided exogenously (Figure 2), resulting in wire-like superstructures in *B. subtilis* and in more spherical flower-like superstructures in *S. oneidensis*. We wondered if these phenotypes remain in the consortium containing dual species. Using the FRET signal from TOTO™-1 / SYTO™60 staining (Figure S5) and NMP staining (Figure S6) indicative of GQ-DNA, we confirmed the presence of GQ-DNA in wires formed after day 3 of dual species biofilm growth in *S. oneidensis* (*tn7*::CFP, in blue) co-cultured with *B. subtilis*. We, furthermore, noticed that the “wires” had CFP signal, suggesting that the CFP from *S. oneidensis* aggregated and integrated into the “wires”. Next, we applied TSA and cDNA labelling to ultimately validate the function and the sequence of the RCR products within those “wires”.

As we cannot exclude natural peroxidase activity present in such giant wire-like supramolecular systems, we recruited an alternative template T2, resulting in a RCR product that is complementary to the cDNA probe but does not form GQ (Table 1). Figures 6A-C show examples of the DNA “wires” detected in the 3-day biofilms of the *B. subtilis* + *S. oneidensis* with the original T1-template (A), an alternative T2-template (B) or without template (C). As we hypothesized, the tyramide signal was detected only in the wires, in which GQ-DNA was synthesized from T1 template (due to either the DNAzyme activity of GQ/ hemin or GQ scaffolding cytochromes and other alternative redox molecules). No tyramide signal was found in the wires enriched with R-DNA synthesized from T2 template.

**Figure 6.**
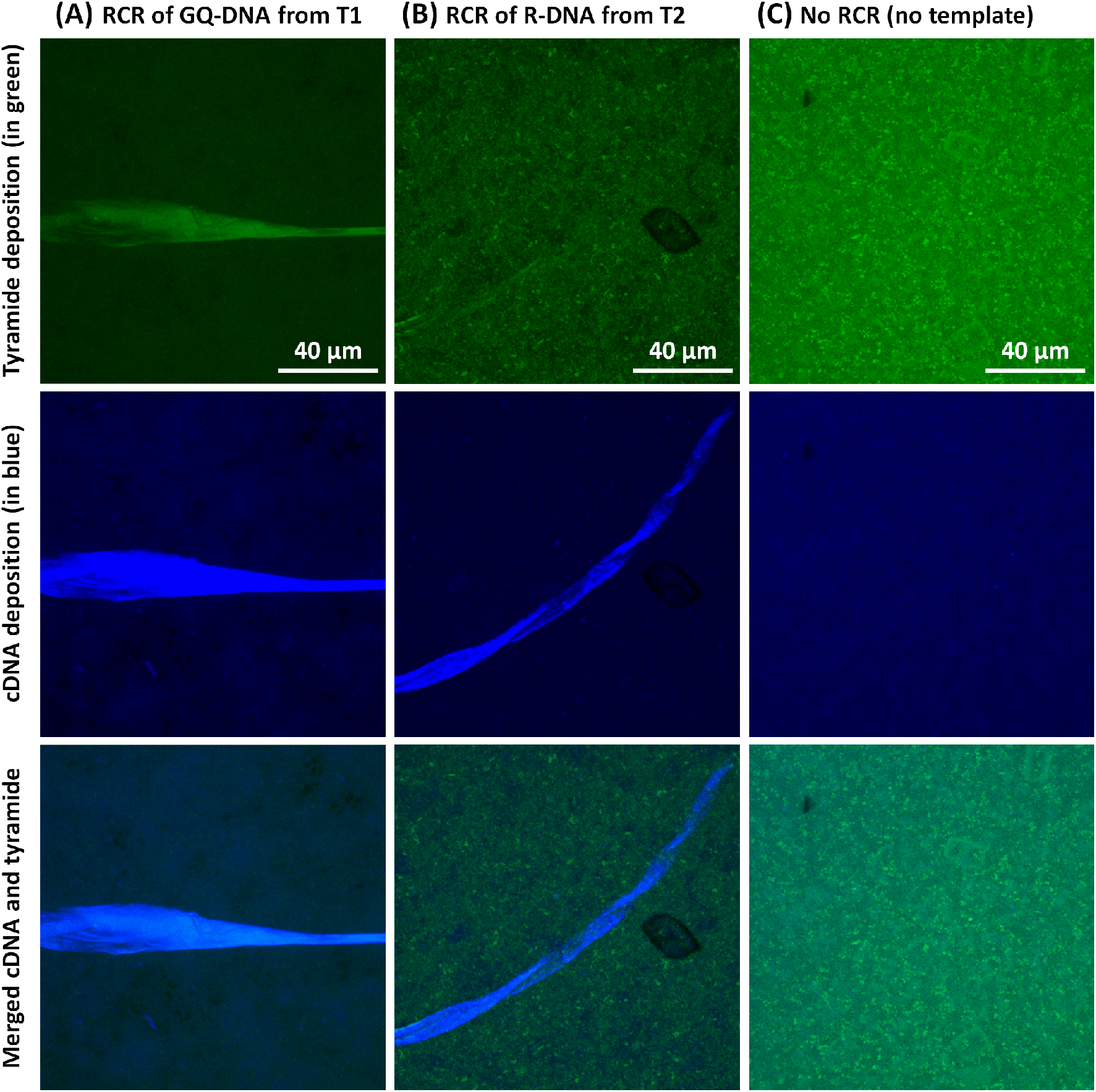
The synthesis of DNA from circular DNA templates in co-cultured *B. subtilis + S. oneidensis* leads to formation of crystal-like “wires” with distinct catalytic function. CLSM images of the wire-like superstructures labelled using elaborated tyramide signal amplification and cDNA labelling. Biofilms were doped with (A) 150 nM circular DNA template T1 for GQ-DNA and (B) 150 nM circular DNA template T2 for R-DNA production. (C) Biofilms were grown without addition of any template. The 3-day biofilms were stained by tyramide residues (Alexa488) in the presence of 0.5 mM ATP and 0.2 % H_2_O_2_ following addition of 5 μM cDNA (5’-Alexa647). See Figure 5 for the assay illustrations. (A), (B) and (C) refer to three locations in three distinct samples.

We hypothesized that the wire-like superstructures are formed from a DNA concatemer produced by RCR from each of the circular DNA templates, which were added during biofilm formation, and that the replication requires activation of prophages in both *S. oneidensis* and *B. subtilis*. No spherical structures were detected in the dual-species biofilms, suggesting a leading role of *B. subtilis* in multispecies biofilm, which also agrees with the faster rate of lytic RCR detected in Figure 2. The replication machinery of bacteriophages in *B. subtilis* may be superior to replication of phage λSo in *S. oneidensis*. The carbon-rich composition of the resulting “wires” implies that such superstructures are formed in response to cellular lysis, like we propose in Figure 2. In the dual-species biofilms of *B. subtilis* + *S. oneidensis* S2391 reporter strain (*tn7*::eCFP, *cro*::Venus), we detected the same featured superstructures (i.e. the “wires”) containing fluorescent protein CFP (Figures S5 and S6). Therefore, the superstructures include self-assembly of GQ-DNA with other biopolymers, and these originate from both microorganisms.

We demonstrated detection of the GQ superstructures within the “wires” in all T1 templated *B. subtilis* biofilm samples using the FRET pair of dyes TOTO™-1 and SYTO™60 (Figure S5), the GQ-specific dye NMP (Figure S6), the complementary cDNA probe and ultimately the TSA (Figure 6A). Furthermore, we demonstrated detection of R-DNA in the T2 templated biofilms using the same complementary cDNA probe (Figure 6B). Therefore, we used T2 as control for synthesis of DNA “wires” that are poor in GQ content. We therefore propose that our approach to DNA synthesis is generic and is programmed by the circular DNA template that we designed for replication of GQ-DNA and random (non-GQ) R-DNA (Table 1), both resulting in formation of the “wires”. However, only GQ-rich DNA “wires” had the catalytic peroxidase-like activity detected by TSA (Figures 5 and 6). Our findings suggest that the synthesis of the superstructures is controlled via sequence design of a small circular template given to both *S. oneidensis* and *B. subtilis*.

## DISCUSSION

We demonstrate for the first time that bacterial prophages can be exploited for programmable synthesis of functional extracellular DNA superstructures through template-directed rolling circle replication. Our real-time monitoring reveals that DNA synthesis occurs predominantly in the extracellular environment (Figures 1 and 2), coinciding with prophage-induced cell lysis (Figure 1). While previous studies of eDNA production in biofilms focus on how eDNA is released from bacterial cells (44), we now show that eDNA can be synthesized from nucleotides directly in the extracellular matrix by phage polymerases. This has implications for how eDNA might be organized in the extracellular matrix, as formation of ssDNA via this mechanism can impact DNA crosslinking in the biofilm, formation of non-canonical DNA structures, DNA-protein interactions and other implications of eDNA in biofilms. Hence, our findings reveal an overlooked dimension of phage-bacteria interactions where prophages contribute to biofilm architecture through extracellular DNA synthesis.

Hundreds of kilobases are generated using rolling circle replication (RCR) from a circular DNA template by a robust phage polymerase, e.g. phi29, in a controlled cell-free environment (30). By designing the circular template, the DNA products of RCR are programmed to fold into a desired conformation, e.g. G-quadruplex (GQ). GQs are highly in demand as these structures can bind a plethora of molecules including redox-active hemin (45), riboflavin (39), cytochrome (46) etc., yielding modular functional electrocatalytic networks (47) and potential for hierarchical assembly. Multiple GQs were shown to assemble intermolecularly resulting in 10–400 nm wire-like nanostructures of ≈2 nm in height and ≈20 nm in width (48). These characteristics make them ideal as building blocks to assemble complex superstructures and further organize various bio- and inorganic molecules. GQ wires containing a large number of stacked guanine tetrads can provide a more efficient π-stacking compared to B-DNA (49), yielding a scaffold and ionic pathways for biobatteries (50). Furthermore, GQ-DNAzymes with peroxidase activity were demonstrated to break graphene oxide (7), suggesting there is an overlooked power of DNA to be harvested.

We first designed a template for synthesis of GQ-DNA because GQ-DNA is resistant to nucleases, and can be differentiated from other DNA structures by immunolabelling (4) and GQ-specific DNA-binding dye NMP (33). Moreover, we used a combination of cell-permeable SYTO™60 and impermeable TOTO™-1 FRET pair (38) to differentiate the extracellular and intracellular GQs (Figure S2). Thus, our template-directed synthesis of GQ-DNA was designed to investigate if such specific DNA sequence could be synthesized extracellularly, but it also provides a new tool for assessing phage-mediated DNA synthesis in bacterial cultures and biofilms in real-time. GQ-DNA is easily detected by fluorescence, and its production requires prophage-triggered release of a polymerase into the extracellular space. The T1-templated RCR assay could therefore be used as an elegant method to study the dynamics of prophage activation in bacteria more broadly, although limited to the prophages that use rolling circle replication.

The GQ-DNA synthesis coincided with *S. oneidensis* lysis and simultaneous phage proliferation, resulting in ≈120 min delay in the DNA synthesis in the cultures doped with mitomycin C and in ≈300 min delay in starved cultures of *S. oneidensis*. In the former case, the timing agrees well with Thöneböhn *et al*. reporting 100 min lag phase before phage assembly and cellular burst (15). Thus, short-term starvation is likely to induce the phage proliferation at the prolonged lag phase, and this could be a generic and sustainable way to activate prophage rolling circle replication.

We prompted the synthesis of GQ-DNA concatemers that folded into superstructures with remarkable morphological diversity that appears to be species-dependent. In the two scenarios, the shape of the DNA is forming after 240 min of incubation under nutrient-limiting conditions (Figure 2). The resulting shapes, which are spherical for *S. oneidensis* and wire-like for *B. subtilis*, can be explained in terms of self-assembly of DNA with lipids and other macromolecules leading to formation of various amphiphilic complexes (51). The “wires” are thought to be bundles of coiled DNA, phospholipid and peptidoglycan strands derived from Gram-positive *B. subtilis* bacteria.

Extracellular DNA (eDNA) plays a predominant role in proliferation of various microorganisms including genus Shewanella and Bacillus (52). We employed *S. oneidensis* and *B. subtilis* as factories of unlimited synthesis of extracellular DNA under nutrient-limiting conditions. *S. oneidensis* is known for its great potential in various biotechnological applications ranging from environmental bioremediation, metal recovery, and material synthesis to bioenergy generation (53). Furthermore, *B. subtilis* is being harnessed for several applications such as microbial fuel cells (54) and even a battery (55). However, both species produce biofilms that are poor in extracellular DNA, where eDNA is predominantly existing as dead cells. Efforts to encapsulate *S. oneidensis* into layers of DNA give eDNA a new role to facilitate electric current generated by *S. oneidensis*, suggesting eDNA may be a key to construct microbial fuel cells (8).

Microbial fuel cells or electromicrobiological processes (e.g. electromethanogenesis for CO_2_ capture) are often limited by the electron transfer processes between the electrode and the associated microorganisms. Therefore, new approaches to harness extracellular nucleic acids for building biofilms are being investigated. For example, enriching environmental biofilms with GQ-DNA can be a way to scaffold bacterial redox-active molecules such as cytochromes, maintaining them in spatial arrangement for a rapid electron transfer (3). Moreover, GQ-rich eRNA is suggested to drive electromethanogenesis in a methanogenic archaeon (6). Thanks to the formation of DNA superstructures in *S. oneidensis* biofilms, we detected an increased level in current in the DPV analysis (Figure 3) suggesting enhanced redox active processes. Thus, the template-programmed GQ-DNA superstructures showing enhanced electrochemical activity in *S. oneidensis* biofilms suggest applications in bioelectronics (56,57), biocatalysis (7,58) and environmental biotechnology (59).

While our results clearly demonstrate prophage-mediated DNA synthesis, several mechanistic questions remain unresolved. Further analysis is required to understand organization of various biopolymers within the superstructures as their formation is most likely dominated by the lipids, proteins, polysaccharides and other macromolecules (in addition to DNA). This may result in inhibition of RCR and the formation of the “empty” spherical and wire-like composites (i.e. without the DNA templates and RCR products). Furthermore, the interplay between various macromolecules within the superstructures may reduce the electrocatalytic properties of GQ-DNA by either mis-folding these structures or shielding it from the electron acceptors. These concerns can be addressed by addition of special enzymes that can selectively cleave each of the macromolecules that are entangled in the superstructures.

### Conclusion

To our knowledge, this is the first time rolling circle replication has been orchestrated in complex multicellular environment, where synthesis of DNA may be slow but unlimited in resources. Thus, this work establishes prophage-mediated rolling circle replication as a programmable tool for engineering programmable catalytic DNA architectures in living biofilms. *S. oneidensis* and *B. subtilis* are just two examples of environmental genus, in which phage-controlled replication of DNA resulted in self-assembly of DNA “flowers” and giant DNA “wires”. However, the incidence of phages that replicate via rolling circle replication is not rare. Future studies should address how this capability can be harnessed to modulate biofilms for biotechnological applications, or to simply synthesize and purify DNA superstructures for applications in biodegradation and tackling environmental challenges.

## MATERIALS and METHODS

### DNA oligonucleotides

### Synthetic DNA ligation

DNA universal primer (5 μM) and one of the linear templates T1 or T2 (4 μM) were gently mixed in 1X Ampligase buffer (from BioSearch) and Ampligase (150 u/mL from BioSearch). The circularization of DNA templates T1 and T2 (Table 1) was obtained by annealing of 5’-phosphorilated single-stranded DNA “padlock probes” onto complementary DNA target at 60°C (20 min), 55°C (20 min), and 50°C (20 min) and simultaneous ligation using 150 u/mL Ampligase (Biosearch) (30). Subsequently, the ligated DNA cooled down gradually to room temperature for another hour. The ligated DNA were applied for rolling circle replication of multimeric GQ and random DNA from the two respective DNA templates (Table 1).

### Synthetic rolling circle replication

Synthesis of GQ-DNA was achieved via rolling circle replication from template T1 using phi29 polymerase (30). Reagents: DNA circle (4 μM), phi29 polymerase (10 u/μL, Thermofisher), phi29 buffer (10×, Thermofisher), BSA (20 mg/mL, Merck), dNTPs (100 mM, NEB), N-Methyl Protoporphyrin (250 μM, NMP, LivChem). A set of 220 μM dATP, dTTP, dCTP and 880 μM dGTP was mixed with 12.5 nM, 50 nM and 100 nM DNA circle in 1X phi29 buffer (33 mM Tris-acetate, 10 mM magnesium acetate, 66 mM potassium acetate, 0.1% Tween 20, 0.2 mg/mL BSA, 1 mM DTT, pH 7.9) in MilliQ water. The mixture was split into 3×3 array: (1) three samples containing three DNA concentrations and phi29 polymerase, (2) three samples containing phi29 polymerase but no DNA, (3) three samples containing three DNA concentrations and no phi29 polymerase. RCR was triggered by adding phi29 polymerase to the groups (1)+(2) and all samples were placed into CLARIOstar plate reader at 37°C. The excitation/emission was 405/615 nm for NMP.

### Bacterial cultures used for prophage induction in planktonic cultures and biofilms

To construct a reporter for LambdaSo capsid production in *S. oneidensis* MR-1, we integrated a second copy of the capsid-encoding gene SO_2963 fused to *sfgfp* into the LambdaSo chromosome by sequential crossover. To avoid subsequent homologous recombination between the two SO_2963 genes and loss of one of the copies, we used a synthetic version of the gene (SO_2963EC). This version was synthesized by Genscript (Rijswijk, NL) and was codon-optimized for E. coli (see sequence below, delivered in pUC57 (Thermo Fisher Scientific), resulting in a gene with 75% DNA sequence identity to the original LambdaSo gene.

To construct the hybrid gene by Gibson assembly (60) the appropriate DNA fragments were amplified by PCR using the given primers (Table 2).

**Table 2.**
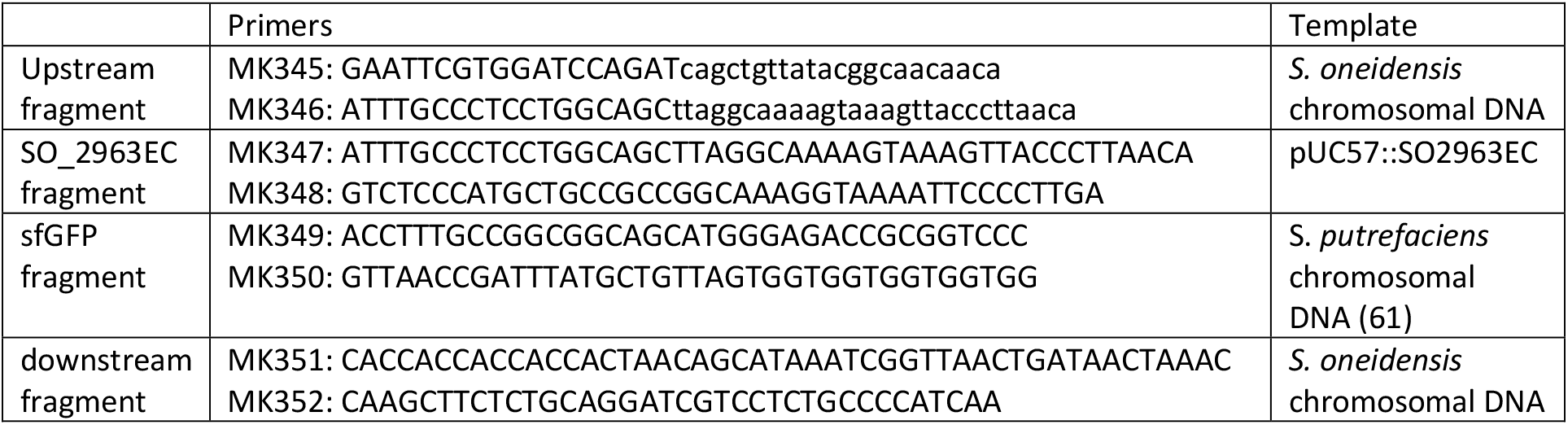
Primers for constructing hybrid gene SO_2963 fused to *sfGFP* in *S. oneidensis* S5558.

The PCR reactions contained HF buffer (5x; Fisher Scientific; CatNo F518L), dNTP mix (stock concentration 10 mM each; Roth; CatNo K039.1), primers (Microsynth; 50 μM each), Phusion polymerase (2U; Fisher Scientific; CatNo F530L) and 50 ng template DNA. Denaturation was carried out at 95 °C, annealing was done at 57 °C, elongation at 72 °C, each for 30 seconds. The cycle was repeated 30 times.

The obtained fragments were gel-purified (NucleoSpinTM Gel and PCR Clean-up Kit; Fisher Scientific; CatNo 11992242) and then fused with each other and EcoRV-digested pNTPS138-R6K (62) (EcoRV obtained from NEB; CatNo #R3195). The reaction contained HiFi DNA Assembly Master Mix (5x; NEB; CatNo E2621L), 50 ng vector DNA and about 150 - 300 ng PCR derived fragments. The mixture was incubated 1h at 50 °C and then directly used for transformation into *E. coli* DH5alpha-LambdaPir (63). The insert of the resultant plasmid was sequenced by SeqLab for confirmation using standard M13 primers. The correct vector was then transformed into the strain *E. coli* WM3064 (W. Metcalf, Urbana, Champagne), from which the plasmid was conjugated into the recipient *S. oneidensis*. Selection for primary and secondary crossover was carried out as previously described (62). Candidate *S. oneidensis* colonies were checked by fluorescence microscopy and PCR. The resultant strain S5558 produced active LambdaSo phage particles and displayed green fluorescence upon production of the capsid.

Sequence of SO_2963EC: ATGCCGAACCCGAACTTTGAAAAACAAATAGAGGAGTTAGGCGTTAACTTGACCAAGATCGGTGACCAGATTAAGAGCGCCGCAGAGGAAACCAATAAGCAGATAAAAGCCTCAGGCGAAATGCATGCTGAAACCAGAGATAAGGTAGACAAGTTGCTGAGCGAACAAGGTGCGTTACAAGCACGTCTTCAAGAGGCGGAACAGAAGCTGTTAAAAGGAGCGCAGTCCAACCAGCAGGAGAGAGAGCAGTCGATCGGCGAACGGGTCGTTAACGACAAGGAGATGGAGGGAGTTTCATCATCCTTCCGTGGGAGTCGTCGTGTGGGAATGCCGCGTAGTGCGATAACAAGCGCGACAGGCAGTGGCGCGGGCCTGGTCCGGCCCGATCGGATGGCGGGAATAGTTGCTGGACCGGAACGTCGTCTTACCATTCGTGACCTGATCGCACCCGGGGAGACTGAATCGAATACCATCGAGTACGTCAAAGAGACCGGCTTTACTAACAACGCGGCGCCCGTGGCCGAAGGGGCCGCCAAACCTTATTCTGAGATCACTTTTGGGTTAGTATCCGCTGGTGTCCGCACAATAGCTCACTTGTTCAAAGCCTCCCGCCAGATTCTTGATGATAGCAAACAATTACAGAGTTTCATAAATGCCCGTGCCCGTTATGGGTTGATGCTGAAGGAGGAAGTTCAGCTTCTTTACGGTAATAATACAGGAGCGAATCTGCACGGAATTATGTTACAAGCTAGC GCCTACGTCAAACCGACAGGGGCGACTGTAGACACTGAGCAGCACATTGACCGCATCCGTCTTGCATTGTTACAGGCAGCCTTAGCTGAGTACGCAGCAGATGGAATAGTATTAAACCCAATCGACTGGGCGGTAATAGAAACTTTGAAAGACTCGAATAAGAATTATCTTATTGGTAAACCCCAAGGGCAAACCGTCCCAACTTTATGGAATCGGCCAGTAGTCGAAACACAGTCAATTGTCCAAAATGAGTTTTTGGTCGGCGCGTTTCAGATGGGTGCGCAAATATACGACCGCATGGATATAGAAGTACTGATTTCAACTGAAAATGACAAAGACTTTGAAAATAATATGGTATCAATACGTGCTGAAGAGCGCCTTGCTCTGGCTGTGTACCGTCCGGAGGCTTTCGTCAAGGGGAATTTTACCTTTGCCTAA

Strain *B. subtilis* (WT) was acquired from DSMZ culture collection (DSM-10). *S. oneidensis* and *B. subtilis* strains were streaked onto LB agar plates and incubated for 24 hours at room temperature and 24 hours at 30°C. A single colony was transferred to LB media to obtain overnight cultures at 30°C reaching stationary/starved phase (OD ≈2.5). Biofilms of *S. oneidensis* were grown in TSB media with 100 mM NaCl or KCl and optionally 20 μM iron sulphate or 20 μM mitomycin C (to test prophage induction).

A buffer was developed for prophage induction in planktonic bacteria: 0.5x phi29 buffer (16.5 mM Tris-acetate, 2 mM (CH_3_COO)_2_Mg, 33 mM CH_3_COOK, 0.05% Tween 20, 0.5 mM DTT, pH 7.9) containing optionally FeSO_4_ (20 μM) or MMC (20 μM) in milliQ. Bacterial cells were harvested from overnight culture, resuspended to OD 5 in the buffer and incubated in Ibidi 96-well plate for 24 h at 30°C to induce prophage-associated rolling circle replication. The buffer was subsequently replaced with TSB media (with 100 mM NaCl or KCl), and the biofilms were grown for 48 hours with and without exogenous 20-200 nM circular DNA template.

To boost possible intracellular replication, a “heat-shock” step was introduced, in which bacterial cells were pre-incubated at 42°C (2 min) and 5°C (1 hour) in the working buffer, prior to the step of prophage activation and replication. This led to crystallization processes inside deformed cells of *S. oneidensis* and formation of larger crystal shapes, which were very distinct from spherical superstructures.

### Fluorescence microscopy imaging of biofilms

Fluorescence microscopy was conducted using two confocal microscopes - the single-photon counting Luminosa microscope (PicoQuant) and Zeiss LSM700. Samples were measured in μ-Slide 8 well IbiTreat slides (Ibidi, Cat.No.: 80826) and in 96-well IbiTreat plates (Ibidi, Cat.No: 89626), respectively, at room temperature.

The Luminosa microscope is based on the inverted Olympus IX73 microscope with a 60x PlanApochromat water immersion objective (NA = 1.2) and Single Photon Avalanche Diode (SPAD) detectors. The fluorescence images were scanned in two different sizes: (1) 10×10 μm of 128×128 pixels with a pixel size of 78 nm and pixel dwell time of 20 μs for imaging the uptake of DNA template T2 in planktonic cultures and (2) 200×200 μm of 1024×1024 pixels with a pixel size of 195 nm and pixel dwell time of 5 μs to obtain fluorescence-based imaging of biofilms.

The Zeiss LSM700 microscope is equipped with a 40×/NA1.4 and 63×/NA1.4 Plan-Apochromat objective using 405, 488 and 639 nm excitation (MBS 405/488/555/639). The Zeiss LSM700 images were scanned as 2D as well as 3D in sizes of (1) 160×160 μm and (2) 100×100 μm of 1024×1024 pixels.

### Real-time detection of phage proliferation and rolling circle replication in planktonic cultures

Bacterial cells were harvested from overnight cultures, resuspended to OD 5 in the working buffer (16.5 mM Tris-acetate, 2 mM (CH_3_COO)_2_Mg, 33 mM CH_3_COOK, 0.05% Tween 20, 0.5 mM DTT, pH 7.9) with and without circular T1-template.

We used *B. subtilis* and various *S. oneidensis* reporter strains and 40 μM NMP to measure molecular processes of phage DNA replication and capsid assembly and extracellular synthesis of GQ-DNA, respectively. The harvested cells were transformed into 96-well plate and incubated for 8 h at 30°C in a CLARIOstar plate reader using following excitation/emission settings: 405/615 nm (NMP), 435/485 nm (*S. oneidensis* CFP), 480/520 nm (*S. oneidensis* GFP and Venus protein). To quantify the NMP signal associated with the GQ-DNA synthesis from GQ-DNA template, we used “NMP difference” obtained by subtracting the NMP signal in “planktonic cells ÷ DNA” from the NMP signal in “planktonic cells + DNA”. To quantify the NMP signal associated with the GQ-DNA synthesis by exogeneous phi29 polymerase, the “NMP difference” was obtained by subtracting the NMP signal in “planktonic cells + DNA ÷ phi29” from the NMP signal in “planktonic cells + DNA + phi29”. The fluorescence of CFP, Venus and GFP and the “NMP difference” were plotted vs. time (2 min data pitch).

We used the single photon counting Luminosa microscope to detect an uptake of the T2 template circularized onto the fluorescently labelled universal Alexa405-primer (Table 1) in planktonic cultures of the wild-type *S. oneidensis* and *B. subtilis* strains. DNA location was monitored after 0, 240 and 360 min of continuous incubation of *S. oneidensis* and, separately, *B. subtilis* bacterial cells with DNA. For detection of the lipophilic membrane dye FM 4-64 (Thermofisher) and Alexa405-DNA, the dyes were excited using the 488 nm laser (LDH-D-C-485S, around 0.1-0.5 μW) and 405 nm laser (LDH-D-C-405S, around 0.1-0.5 μW), respectively. Alexa405 was detected around 430-460 nm (BP445/30) whereas FM 4-64 was detected around 663-735 nm (BP700/75). A laser repetition rate of 40 MHz was used. FM 4-64 stained cellular membranes and exogeneous Alexa405-DNA were visualized in red and blue, respectively. The red channel was kept at a maximum of 200-500 counts, and the blue channel was set to be 100x of the red (maximum of 2-5 counts) for images from 0-240 minutes and 10× of the red (maximum of 20-50 counts) for images from 360 minutes.

### End-point detection of rolling circle replication products in biofilms using DNA binding antibodies

Biofilms were immunolabelled in 3 % BSA. After a 5 min blocking step with 3% BSA in 1 × PBS, *in vitro* biofilms in separate microwells were incubated with 60 μL solutions of Atto488-BG4 antibody (10 mg/L, Absolute antibodies) in the 3% BSA for 75 min. Second, we added a 60 μL solution of mouse AB1 antibody (5 mg/L, Abcam) in the 3% BSA and continued incubation of the biofilms with the two antibodies for another 75 min. Third, after washing the biofilms with 120 μL of the 3% BSA, the biofilms were incubated with 60 μL solution of anti-mouse CF405S-AB2 antibody (10 mg/L, Merck) in the 3% BSA. The biofilms were washed in the 3% BSA and, finally, stained in 10 mg/L solution of FM4-64 in 100 mM NaCl. The stained biofilms were visualized by confocal laser scanning microscopy (CLSM, Zeiss LSM700) showing BG4 in green, AB1-AB2 in blue and FM 4-64 in red. The immunolabelled *S. oneidensis* biofilms were visualized by confocal laser scanning microscopy (CLSM, Zeiss LSM700) using 63x oil-immersed objective.

### End-point detection of rolling circle replication products in biofilms using DNA-binding dyes and FRET

DNA-binding dyes N-Methyl Protoporphyrin (250 μM, NMP, LivChem), SYTO™41, SYTO™60, and TOTO™-1 (250 μM, Thermofisher) were applied to biofilms in 100 mM NaCl solution. After biofilm formation, the media was exchanged to 100 mM NaCl solution with NMP and SYTO™41 (for a quick staining) or with TOTO™-1 and SYTO™60 (for FRET staining) just before imaging.

*In vitro* biofilms of *B. subtilis* + *S. oneidensis* S2391 (live cells = CFP, ex. 405 nm, em. 300-550 nm, gain 1000) were incubated with 60 μL solution of 25 μM NMP (ex. 405 nm, em. 600-800 nm, gain 1000) to visualize GQ-DNA. *In vitro* biofilms of *S. oneidensis* (wild-type MR-1) were incubated with 60 μL solution of 25 μM NMP (ex. 405 nm, em. 600-800 nm, gain 1000) and 12.5 μM SYTO™41 (ex. 405 nm, em. 400-600 nm, gain 650) to visualize GQ-DNA and intracellular DNA in the live cells, respectively. GQ-DNA and live cells were presented in red and blue, respectively. *In vitro* biofilms of *S. oneidensis* and *B. subtilis* (both single and multispecies) were incubated with 60 μL solution of 2.5 μM TOTO™-1 (ex. 488 nm, em. 400-630 nm, gain 900) and 12.5 μM SYTO™60 (ex. 639 nm, em. 630-800 nm, gain 900) to visualize total eDNA and live cells, respectively. The eDNA and live cells were presented in green and blue, respectively. Since the two dyes bind GQ-DNA (38), we used FRET (donor ex. 488, acceptor em. 630-800 nm, gain 900) to visualize GQ-DNA, which was presented in red. The biofilms were visualized by confocal laser scanning microscopy (CLSM, Zeiss LSM700) using 63x (for *S. oneidensis* monospecies) and 40x (for *B. subtilis* mono- and multispecies biofilms) oil-immersed objectives.

Additionally, *B. subtilis* + *S. oneidensis* S2391 co-cultured biofilms were visualized by single photon counting confocal fluorescence Microscopy (Luminosa). TOTO™-1 (donor) was excited using the 488 nm laser (LDH-D-C-485S, 40 nW – 9 μW) and detected around 500-540 nm (BP520/35). The FRET signal from SYTO™60 was measured upon excitation with the 488 nm laser and detected around 650-725 nm (BP690/70). The direct SYTO™60 signal (acceptor) was excited using the 640 nm laser (LDH-D-C-640S, around 0.5 nW – 0.1 μW) and detected using the same BP690/70 filter. The 640 nm laser was always set to be 80x lower compared to the 488 nm laser. In several experiments, the *tn7*::CFP signal was also measured using the 405 nm laser (LDH-D-C-405S, around 9 μW) and detected around 430-460 nm (BP445/30). For all pulsed lasers, a repetition rate of 20 MHz was used in pulsed interleaved excitation mode. The donor and FRET channels were kept at the same contrast range (for example 0-50 counts), whereas the acceptor channel had half of this contrast range (for example 0-25 counts) for all shown images. The CFP channel was kept at the same contrast setting as for the acceptor channel.

### End-point detection of rolling circle replication in biofilms using cDNA and tyramide signal amplification

Biofilms were labelled with the cDNA oligonucleotide complementary to the products of DNA rolling circle replication (Table 1). We used universal cDNA (5 μM), unlabeled as well as labelled with 5’-Alexa647, to tag products of rolling circle replication generated from various templates. Additionally, we used tyramide signal amplification (TSA) as a method to specifically detect peroxidase activity of GQ-DNAzymes in the presence of hydrogen peroxide. To detect such activity, the biofilms were obtained in TSB media doped with 20 μM hemin (Sigma, Porcine BioXtra, Cat.No 51280) and 20 μM FeSO_4_. Subsequently, the media was replaced by Tris-Acetate buffer (pH 6) with 0.5 mM ATP, 0.2 % H_2_O_2_, and 100 mM KCl, in which TSA was performed for around 2 hours. The biofilms were visualized by confocal laser scanning microscopy (CLSM, Zeiss LSM700) using 40x oil-immersed objective. The cDNA and the tyramide fluorescence were shown in blue and green, respectively.

### End-point detection of rolling circle replication in biofilms using scanning electron microscopy

Scanning electron microscopy (SEM) analysis was performed on biofilms grown in μ-Slide 8 well IbiTreat slides (Ibidi, Cat.No.: 80826). The biofilms were gently rinsed with deionized water, decanted and dried under vacuum overnight, before sputter coating with a thin layer of platinum. The SEM images were acquired using a TESCAN Clara SEM under high vacuum (pressure <0.04 Pa), an accelerating voltage of 5 keV, a beam current of 100 pA and an Everhart-Thornley (E-T) detector for secondary electrons.

SEM energy-dispersive X-ray spectroscopy (EDX) mapping was performed using a beam current of 3 nA.

### Measurements of redox activity of eDNA superstructures in biofilms

Bioelectrochemical analyses of the monocultured *S. oneidensis* biofilms grown aerobically on screen printed electrodes (SPEs) were carried out on the SPEs (Metrohm DropSens DRP-C110, Metrohm Spain). It had a 4 mm diameter graphite working electrode (WE) with surface area of 0.126 cm^2^, a graphite counter electrode, and Ag pseudo-reference electrode. SPEs were surface sterilized in 70% v/v ethanol, washed thrice in sterile deionized water, air dried under aseptic conditions and exposed to UV treatment for 15 min. Biofilms were grown for 3 days on SPEs in 1 mL glass tubes with butyl rubber caps fused to working electrode portion of SPEs prior to analyses (2). Biofilms were grown 3-days aerobically (30°C, 100 rpm) and additionally 3-days anaerobically (30°C, 100 rpm). After incubation and biofilm formation, SPEs with formed biofilms were immediately connected to a PC-controlled multichannel PalmSens4 potentiostat (PalmSens Netherlands). Differential Pulse Voltammetry (DPV) analyses were carried out using electrode potential spanning from - 0.8 V to 1.5 V and following parameters: 5 s equilibration, 0.01 V step, 0.05 V/ 0.3 s pulse and 5 mV/s scan rate. Two biological replicates of *S. oneidensis* were prepared for each test condition and three technical replicates (measurements) for each sample were recorded and averaged. All analyses were carried out in open laboratory conditions and with no disruption of sample.

## ACKNOWLEDGEMENTS

We thank Prof. Ebbe Sloth Andersen, Aarhus University, for providing access to experimental facilities for CLARIOstar fluorescence measurements. We thank Prof. Thomas William Seviour for fruitful discussions of eDNA formation under stringent conditions in environmental bacteria.

## AUTHOR CONTRIBUTIONS

Gabriel Antonio Minero (**GAM**) developed research concepts and acquired the funding to perform most of the presented work. *Shewanella oneidensis* reporter strains and mutants were provided by Kai Thormann (**KT**). Rikke Louise Meyer (**RLM**) provided access to Zeiss 700 CLSM. Mingdong Dong (**MD**) provided access to SEM. All authors contributed to the design of experiments. DNA ligation and monitoring of rolling circle replication in planktonic cultures and cell-free was performed by **GAM**. CLSM imaging of samples stained by antibodies, DNA-binding dyes, cDNA and TSA were performed by **GAM**. Single photon counting confocal imaging and data analysis were carried out by Line Mørkholt Lund (**LML**) and supervised by Victoria Birkedal (**VB**). SEM imaging was performed by Lasse Hyldgaard Klausen (**LHK**). Electroactivity and DPV analysis of biofilms was measured by Obinna Markraphael Ajunwa (**OMA**). The manuscript was prepared by **GAM** while **KT, LML, VB, RLM** and **MD** contributed to writing.

