## Supplementary Information for "Secret life of prophages: template-directed synthesis of DNA superstructures via prophage activation and rolling circle replication in bacterial biofilms"

Gabriel Antonio S. Minero et al.

**SUPPLEMENTARY INFORMATION (Secret Life of Prophages)**

**Rolling circle replication using commercial polymerase and nucleotide triphosphates**

Rolling Circle Replication (RCR) is a well-established technique to replicate a circular DNA using phage polymerases with strand-displacing activity. Figure S1 shows increase of “NMP difference” in RCR templated by the ligated circular DNA (25-200 nM). In this technique, a “padlock probe” or template (shown in blue), 5’-phosphorilated, is first annealed and ligated onto a single-stranded DNA “primer” using Ampligase or another DNA ligase. In the presence of nucleotide triphosphates (NTPs) and polymerase, the 3’-end of the primer (shown in red) is extended into the growing DNA chain, resulting in release of pyrophosphates. After the full circle of DNA template is completed, polymerase uses its strand displacing activity to keep rolling circle replication until all NTPs are depleted or polymerase is inactivated. We used T1 circular template containing triplets of cytosines to generate triplets of guanines foldable into G-quadruplex GQ-DNA as the end-product of RCR in the presence of commercial phi29 polymerase. In the presence of the synthesized GQ-DNA, the N-Methyl Protoporphyrin (NMP) fluorescence lights up that makes it possible to detect such product of rolling circle replication. We deducted NMP fluorescence of non-templated controls from the samples templated with 25 nM, 100 nM and 200 nM circular DNA and show the dose-response signal saturating in all samples around 120 min suggesting inactivation of the phi29 polymerase.

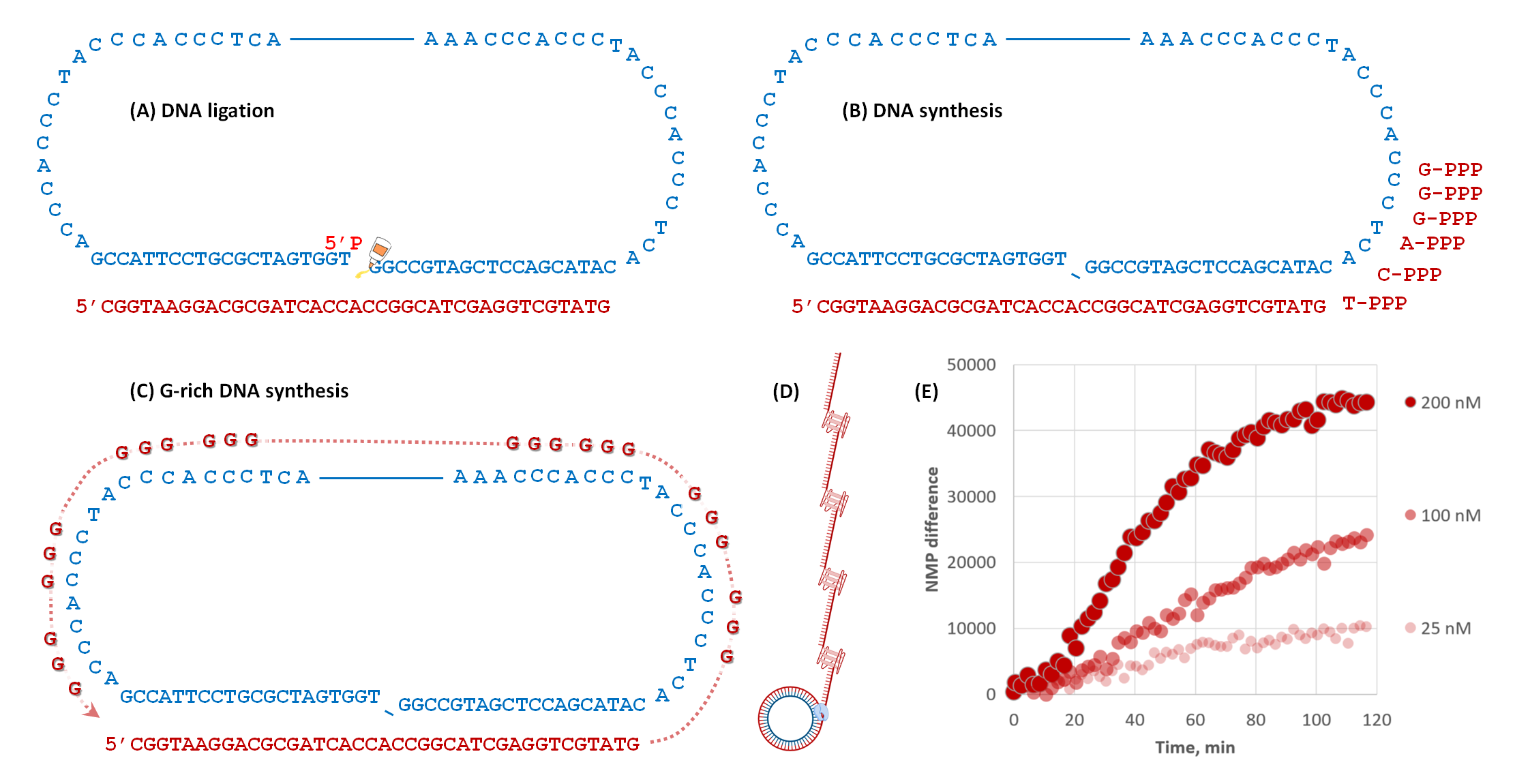

**Figure S1. Real-time detection of rolling circle replication of GQ-DNA from template T1 using phi29 polymerase, dNTP and different concentrations of the circular DNA T1,.** (A) Ligation the circularized DNA template is followed by (B) Incorporation of NTPs, DNA synthesis and, ultimately (C) rolling circle replication resulting in an end-product containing desired sequence, e.g. (D) GQ-motifs (synthesized using T1 template). (E) Detection of the RCR using “NMP difference”.

One turn around T1 template resulted in eight triplets of guanines foldable into two GQs separated by two DNA tracks:

(-**GQ**-TCGGTAAGGACGCGATCACCACCGGCATCGAGGTCGTATGTGA-**GQ**-TTTGTGGCAGGTCAGTCAAGTATACTG CACTATTTGA-**GQ**-TCGGTAAGGACGCGATCACCACCGGCATCGAGGTCGTATGTGA-)n.

**CLSM imaging of T1-templated *S. oneidensis* biofilms using SYTO^TM^60/ TOTO^TM^-1 FRET pair detects GQ-DNA within spherical superstructures**

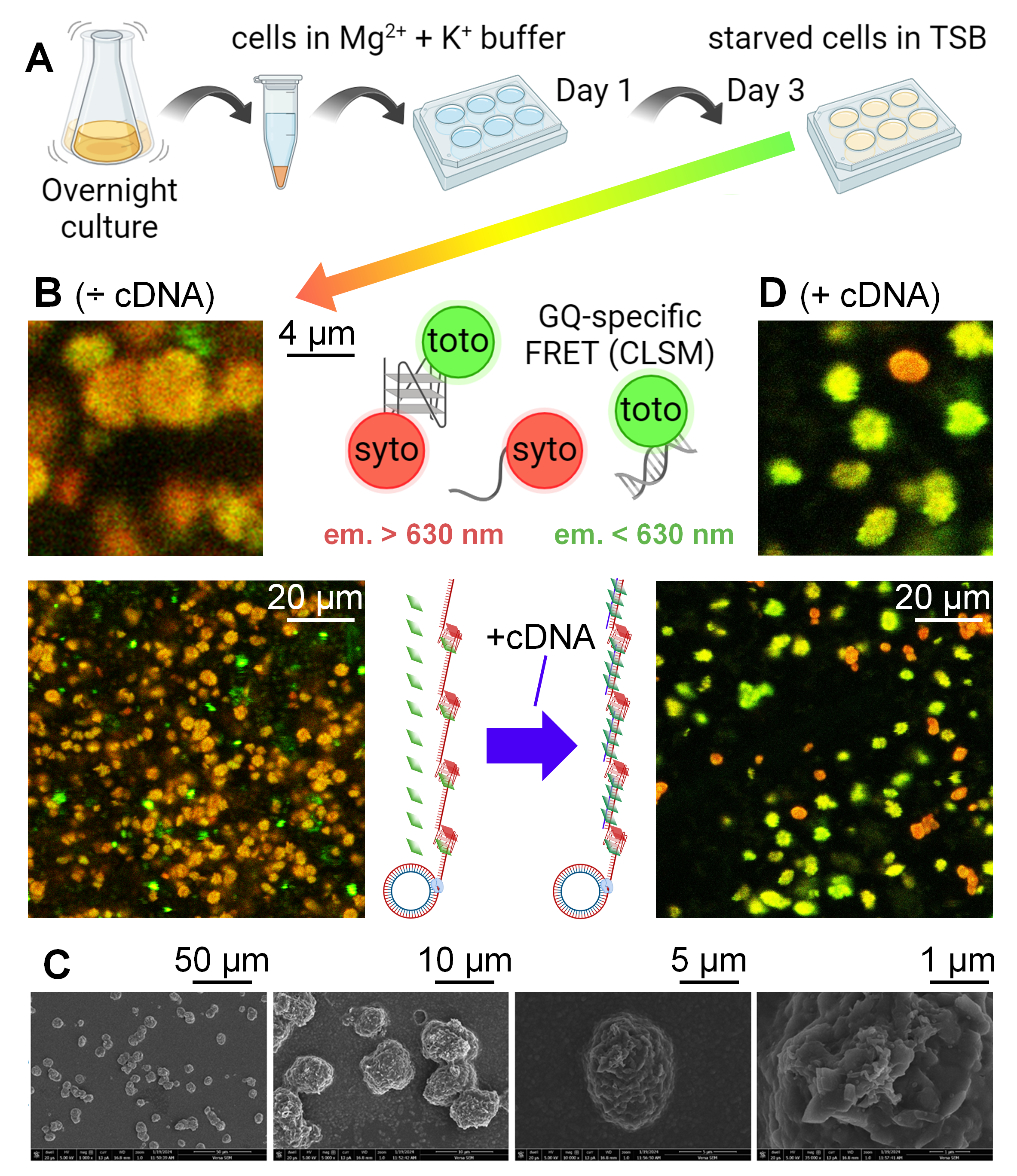

**Figure S2. CLSM of GQ-DNA superstructures replicated from template T1 in 3-day biofilms of wild-type *S. oneidensis* stained with SYTO**^TM^**60/TOTO**^TM^**-1.** (A) The workflow used for biofilm formation with starvation-induced RCR (shown by black arrows). (B) Upon biofilm staining, both TOTO^TM^-1 and SYTO^TM^60 bind to GQ-DNA and cause FRET. High FRET signal (red) detected in spherical structures indicates the end-product of RCR enriched in the multiple GQ motifs. Cartoons of GQ-, single- and double-stranded DNA existing in the extracellular matrix and interacting with the dyes. (C) Replication of the circular DNA T1 in *S. oneidensis* biofilms leads to formation of crystal-like spherical super-structures detected by SEM. (D) The RCR end-product is complementary to cDNA. High TOTO^TM^-1 signal (green) detected in spherical structures indicates an increased amount of dsDNA between cDNA oligonucleotide and the end-product of RCR upon addition of 10 µM cDNA (shown by blue arrow). TOTO^TM^-1 and SYTO^TM^60 are represented by green and red cartoons, respectively.

Figure S2 shows RCR assay first in bacterial cultures under nutrient-limiting conditions (1-day) and then in biofilms grown for 2-days in TSB media. We, furthermore, used live/ dead staining by SYTO^TM^60/ TOTO^TM^-1 DNA-binding dyes to detect synthesized eDNA superstructures. TOTO^TM^-1 normally lights up B-DNA and dead cells (green) and does not light up GQ-DNA. We recently established that GQ-DNA binds SYTO^TM^60 and TOTO^TM^-1 resulting in Förster Resonance Energy Transfer (FRET) (38). To detect GQ-DNA, SYTO^TM^60 (acceptor) emission (> 630 nm) is detected upon TOTO^TM^-1 (donor) excitation (488 nm). To simultaneously detect B-DNA, TOTO^TM^-1 (donor) emission (<630 nm) is detected upon TOTO^TM^-1 (donor) excitation (488 nm). In the extracellular space, we found that TOTO^TM^-1 and SYTO^TM^60 form a FRET pair resulting in SYTO^TM^60 (acceptor) emission (red) upon TOTO^TM^-1 (donor) excitation (Figure S2B). The morphology of the spherical structures forming in the DNA templated *S. oneidensis* WT biofilms was furthermore visualized by SEM (Figure S2C).

Furthermore, we labelled biofilms with cDNA probe (Table 1) forming double-stranded B-DNA with the single-stranded product of RCR to confirm sequence of the RCR product (shown by blue arrow in Figure S2). Using the two DNA-binding dyes allowed us to detect cDNA binding to the DNA products synthesized from template T1. After 1 h incubation of the spherical superstructures (in *S. oneidensis* biofilm, Figure S2B) with the cDNA, a sub-population of the spherical structures increased in TOTO^TM^-1 fluorescence (emission < 630 nm, shown in green), indicating formation of double-stranded B-DNA (Figure S2D). However, another sub-population of spherical structures did not light up TOTO^TM^-1 (did not become more “green”) after cDNA labelling, indicating a barrier to the cDNA probe or the TOTO^TM^-1 dye. These could be, for example, protoplasts with a remaining lipid barrier preventing passage of the cDNA and are distinct from another sub-population of the spherical structures. Therefore, we assumed that there are two phenotypes of biofilms with the RCR induced in (1) the extracellular or (2) the intracellular environment. In contrast to planktonic assay of < 24 h, uptake of the circular DNA templates is, therefore, possible during 3-day biofilm formation.

No FRET as well as no spherical superstructures were detected in the non-templated WT biofilms (Figure S3) or templated *S. oneidensis* MUT (ΔλSo ΔMuSoI ΔMuSoII) biofilms (Figure S4).

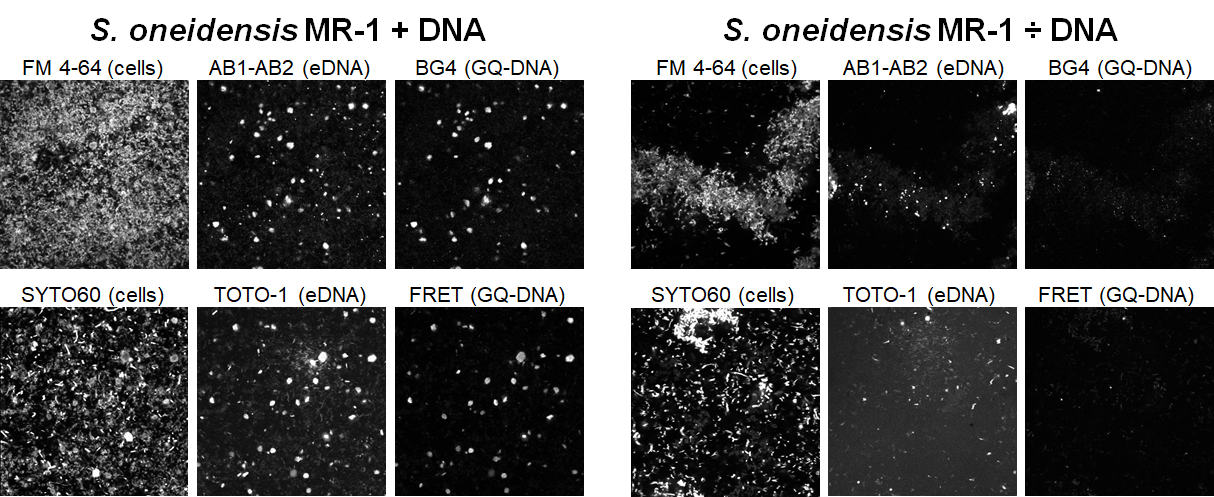

**Figure S3. *S. oneidensis* biofilms doped with circular DNA template T1 form spherical superstructures detected by the GQ-specific immunolabelling and the FRET pair.** (A) Biofilms obtained in DNA-TSB-NaCl containing 200 nM exogenous circular DNA template and (B) biofilms obtained in plain TSB-NaCl were labelled by DNA-specific antibodies BG4 and AB1-AB2 in combination with FM 4-64, as well as by DNA-binding stains TOTO-1 and SYTO60. 2D CLSM 100x100 µm.

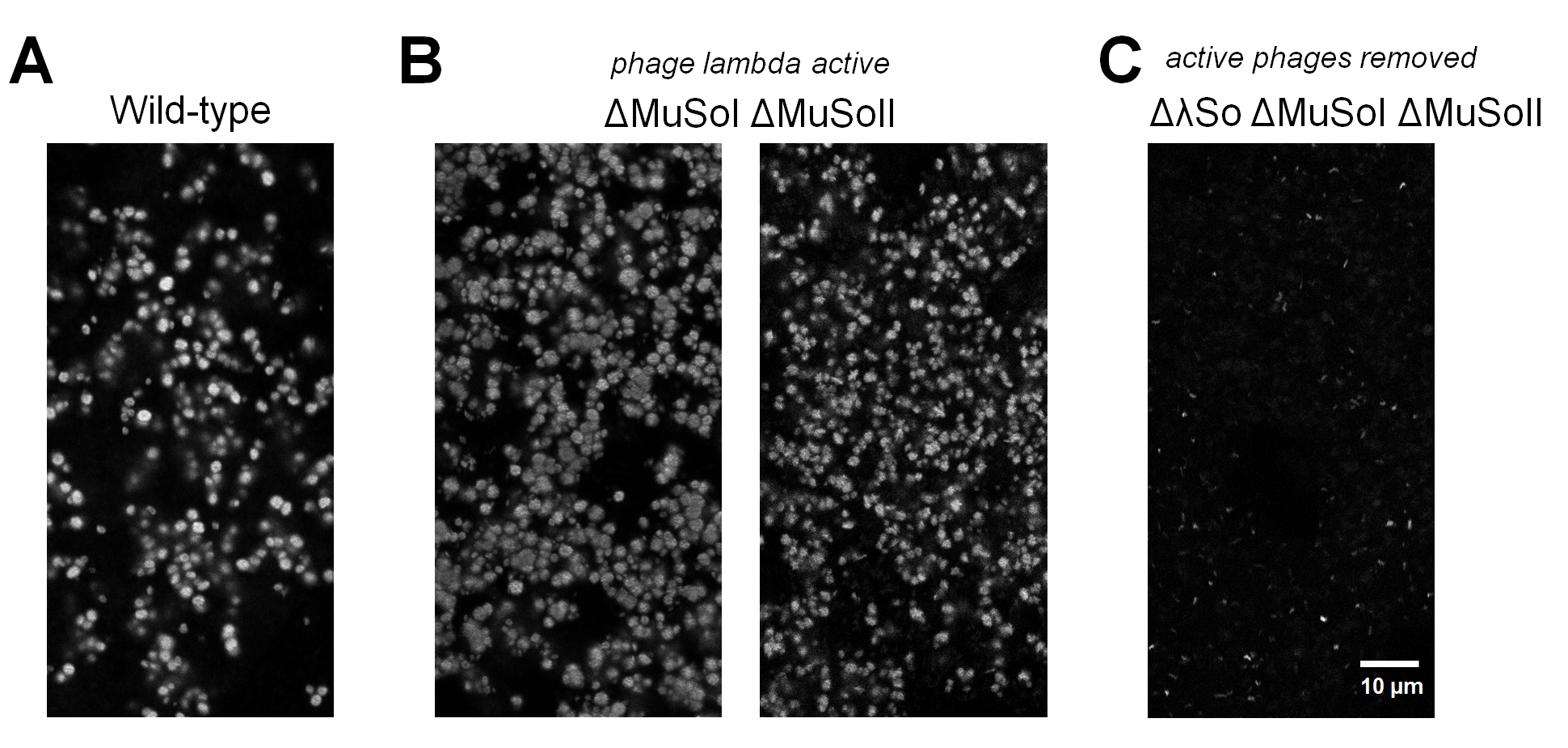

**Figure S4. Prophage λSo proliferation is responsible for extracellular synthesis of GQ-DNA in *S. oneidensis* 3-day old biofilms.** CLSM (FRET channel) of the replicated DNA template T1 in 3-day biofilms of *S. oneidensis* MR-1 strains: (A) WT, (B) ΔMuSoI ΔMuSoII and (C) ΔλSo ΔMuSoI ΔMuSoII stained with TOTO-1 and SYTO60.

**CLSM imaging of T1-templated *S. oneidensis* + *B. subtilis* biofilms using SYTO^TM^60/ TOTO^TM^-1 FRET pair detects GQ-DNA within wire-like superstructures**

Next, we wondered if formation of the wire-like superstructures detected in *B. subtilis* biofilms is influenced by *S. oneidensis* when co-cultured with *B. subtilis.* We used the same assay and workflow as in Figure S2 and the S2391 reporter strain of *S. oneidensis* constituently expressing CFP protein from its tn7 promoter. The biofilms were stained with TOTO^TM^-1 and SYTO^TM^60 couple resulting in FRET signal specific for GQ-DNA. We noticed very high FRET signal (red) from a wire-like structure close to 1 mm in length (Figure S5). Furthermore, we detected the CFP produced from *S. oneidensis* S2391 (*tn7*::CFP) within the wire (Figure S5B) suggesting that CFP among other biopolymers derived from *S. oneidensis* contributes to the wire-like superstructure typically found in *B. subtilis* (in this work).

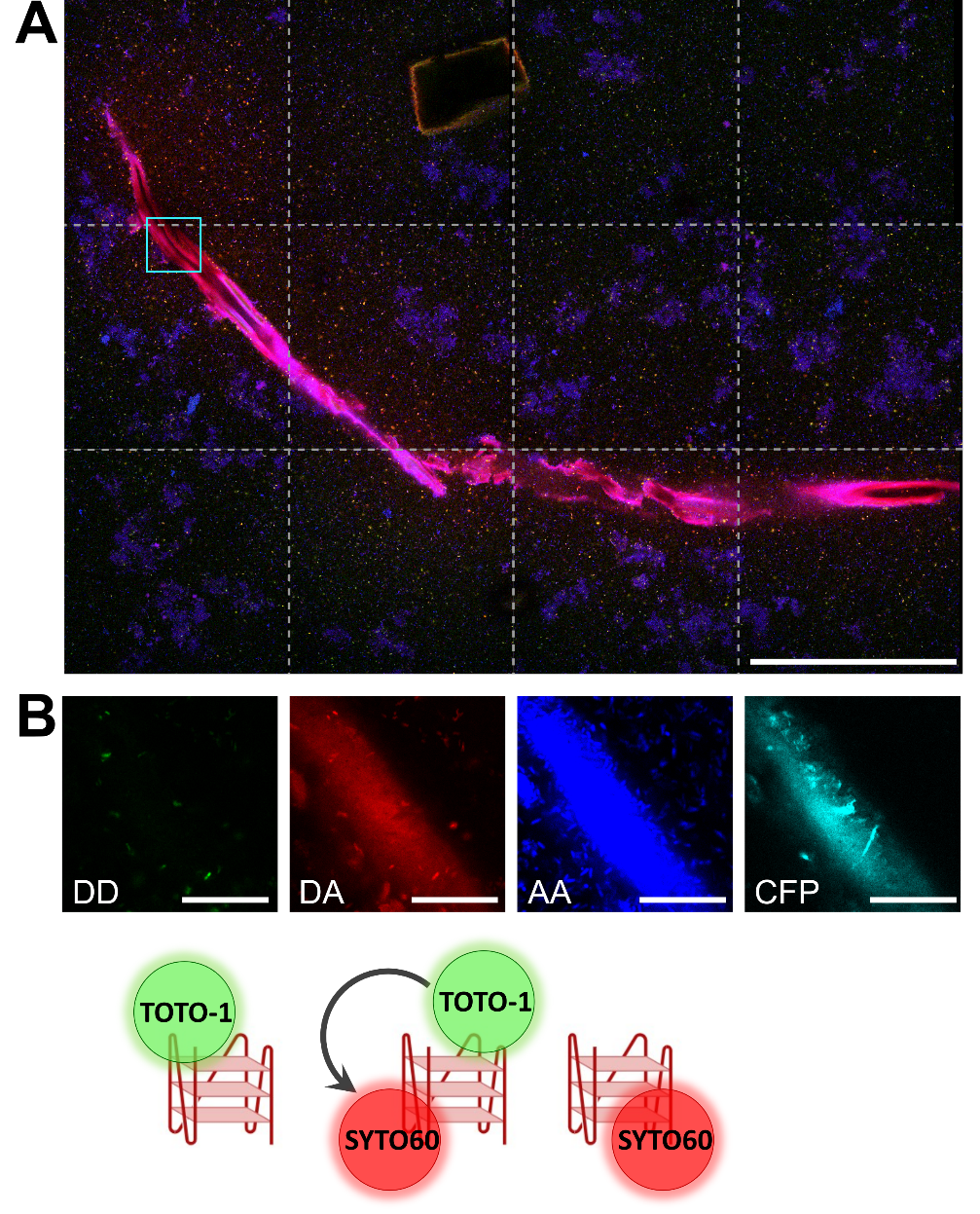

**Figure S5. FRET in GQ-DNA construct produced in *S. oneidensis* and *B. subtilis* (co-cultured) biofilm (day 3) in TSB-KCl detected by single photon counting confocal microscopy.** (A) Example of a millimeter long, twisting wire. Here, green is from the donor (DD, TOTO-1 channel), red is from FRET signal (DA, SYTO60 channel), and blue is from the directly excited acceptor (AA, SYTO60 channel). Since the difference in detection efficiency of green and red signal is 1:2.5, the signal in the green channel has been multiplied by 2.5. Still, the wire is red overall, meaning high in the FRET signal (DA). Scale bar is 200 µm. (B) Small cutout of the same wire in (A), cyan box, at a different z-height to show the surface of the wire. Here, the fluorescent signal is split into four channels. The donor signal in green (DD, TOTO-1 channel), the FRET signal in red (DA, SYTO60 channel), direct acceptor in blue (AA, SYTO60 signal), and *tn7*::CFP signal in cyan (CFP channel). Scale bars are 20 µm.

The CFP expressed specifically by *S. oneidensis* S2391 was found to be present in biofilms of monospecies S2391 as well as co-cultured with *B. subtilis* (Figure S6). Therefore, we believe that the pathways of DNA replication resulting in the giant DNA wires are similar in *B. subtilis* and *S. oneidensis*.

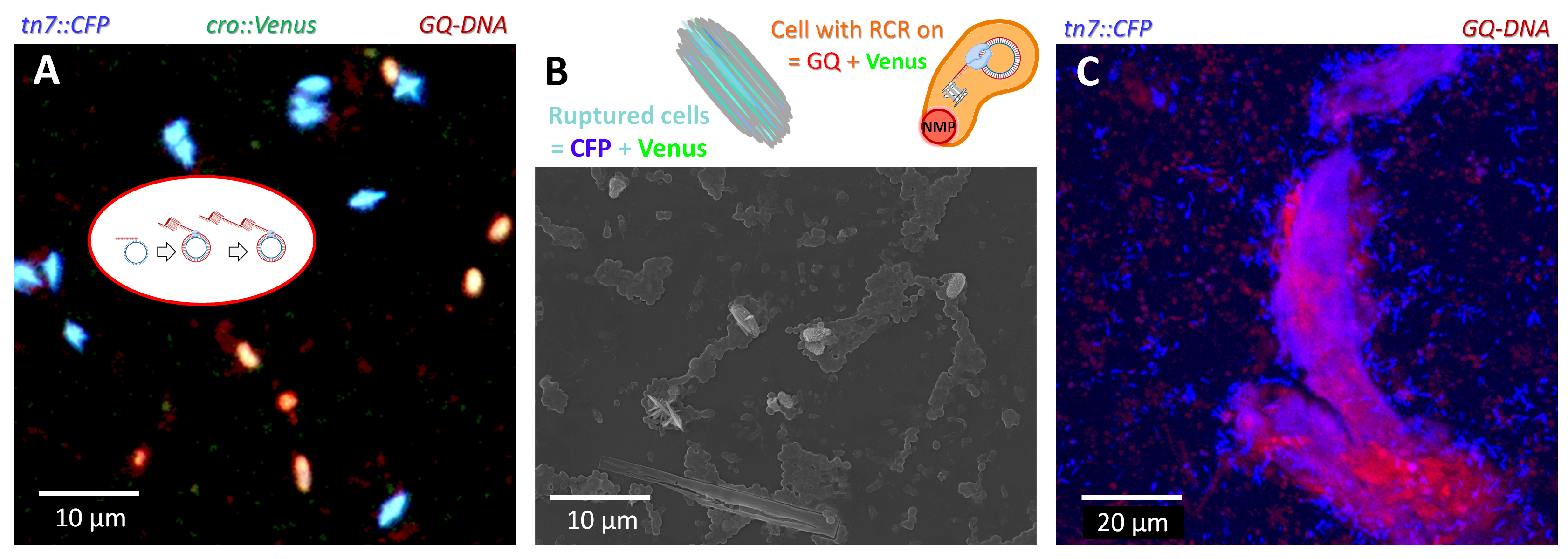

**Figure S6. CFP from *S. oneidensis* is deposited into the supramolecular structures synthesized and self-assembled in 3-day biofilms of *S. oneidensis* S2391 (*cro*::Venus, *tn7*::CFP) co-cultured with *B. subtilis*.** (A) CLSM and (B) SEM images of cells being ruptured. NMP signal predominantly detected inside deformed cells of *S. oneidensis* with active synthesis of *cro* (*cro*::GFP indicative of prophage λSo proliferation, green). Ruptured cells show high signal of both CFP (blue) and Venus (green) proteins. (C) The wire-like superstructure detected by NMP (indicative of the GQ end-product of RCR) and CFP (indicative of *S. oneidensis*) signals.
